# PARG activity is required for cell death by parthanatos

**DOI:** 10.64898/2026.05.12.724507

**Authors:** Rafael Dias de Moura, Beatriz Kopel, Priscilla Doria de Mattos, Penélope Ferreira Valente, Aline Costa Santos, Rafael Teixeira do Nascimento, Fernanda Manso Prado, Marisa Helena Gennari de Medeiros, João Carlos Setubal, Nadja Cristhina de Souza Pinto, Nícolas Carlos Hoch

## Abstract

Cell death by a non-apoptotic pathway termed parthanatos is induced by hyperactivation of the DNA damage sensor PARP1, which uses NAD^+^ as a substrate to catalyse the poly-ADP-ribosylation (PARylation) of proteins. Parthanatos has been implicated in several pathological conditions such as ischaemia-reperfusion injury and neurodegenerative processes, including Alzheimeŕs and Parkinsońs diseases, as well as rare genetic disorders characterized by PARP1 hyperactivation. However, the precise sequence of molecular events by which excessive PARP1 activity causes cell death is currently unclear. Here we show that, in addition to PARP1-dependent PARylation, the execution of parthanatos also requires the hydrolysis of poly-ADP-ribose (PAR) chains by the glycohydrolase PARG. While complete inhibition of PARG activity prevents parthanatos, low levels of residual PARG activity are sufficient to support cell death by this pathway. Due to a phenotypic discrepancy between CRISPR/Cas9-generated PARG KO cells and PARG inhibitor-treated cells, we uncovered a new PARG splice variant predicted to encode an isoform of 53 kDa, termed PARG53, and provide evidence for the incorrect annotation of the previously reported isoforms PARG55 and PARG60. PARP1 hyperactivation during parthanatos leads to rapid and profound depletion of NAD^+^ and ATP pools, but full PARG inhibition only prevents the depletion of ATP, indicating that the depletion of these core metabolites can be uncoupled, and that ATP depletion is more closely correlated with cell death. This work sheds light on the initial steps of parthanatos induction, demonstrating that PAR formation and PAR hydrolysis are both required for this type of cell death.

## Introduction

ADP-ribosylation is a covalent modification of proteins and nucleic acids, which regulates diverse cellular processes such as DNA damage signalling, DNA replication, transcription, and immune responses ^[1–4]^. ADP-ribosyl transferases (ART) enzymes, which include the human PARP family with its 17 members, catalyse the transfer of the ADP-ribosyl group from NAD^+^ to a target biomolecule, with the release of nicotinamide ^[2,5]^. This modification is found as single ADP-ribose units (termed mono-ADP-ribose, or MAR) or as chains composed of hundreds of ADPr monomers (termed poly-ADP-ribose, or PAR). In human cells, PARP1 and PARP2 are activated by single and double strand breaks in DNA, and thus PARylate (i.e., add PAR chains to) chromatin proteins, mainly themselves and histones ^[1–3,5]^. These polymers, which are strongly negatively charged and often branched, lead to a more relaxed chromatin state and act as a scaffold for the recruitment of DNA repair proteins and chromatin remodellers ^[1–3]^. PAR also plays important roles in DNA replication, such as stabilizing replication forks ^[1]^ and regulating Okazaki fragment processing ^[6,7]^. Importantly, PARylation is a transient and dynamic modification, being degraded within minutes by the highly active poly-ADP-ribose glycohydrolase (PARG) enzyme ^[8]^.

PARG is a glycohydrolase of the macrodomain family, with five isoforms characterized at the protein level, which result from alternative splicing in the 5’ end of its transcript. The canonical splice isoform is composed of 18 exons and encodes a nuclear protein of 111 kDa ^[9]^, termed PARG111. Two slightly smaller splice isoforms lack the first and second exons, respectively, and encode the cytoplasmic proteins PARG102 and PARG99 ^[10]^. All three of these isoforms are known to be catalytically active, given that the PARG catalytic domain is located at the C-terminal end of the protein. Two additional isoforms, PARG60 and PARG55, have been reported ^[11]^, but are not thought to be catalytically active. The transcripts reported to encode PARG55 and PARG60 are spliced from exon 1 to exon 4 (skipping exons 2 and 3), as well as from exon 4 to exon 6 (skipping exon 5), but use slightly different e1-e4 splice junctions, leading to different translation start sites ^[8,10–12]^.

Mechanistically, PARG cleaves the glycosidic bonds between ADP-ribose moieties, acting mainly as an exoglycohydrolase that produces free ADP-ribose monomers, but also has endoglycohydrolase activity, producing free PAR ^[13–15]^. Importantly, PARG is incapable of removing the last ADP-ribose monomer from most amino acid sidechains, including serine residues which are thought to be the main PAR acceptor sites after DNA damage ^[15–17]^. Therefore, additional ADPr hydrolase activity provided by enzymes such as ARH3, which can cleave this serine-ADPr bond, is required to completely remove PAR chains ^[18,19]^. PARG activity is thought to proceed much faster than ARH3 activity, generating a wave of PARylation followed by a wave of MARylation, which could have different biological functions ^[8,20]^. Recently, PARG has been shown to hydrolyse mono-ADPr attached to glutamate and aspartate residues, indicating that Glu/Asp-linked PAR chains may be completely reversed by PARG alone ^[16,21]^.

The extent of PARP1/2-dependent chromatin PARylation, which is likely directly correlated to the amount of genomic DNA breaks, can be a determinant of cell fate ^[22]^. While PARylation promotes DNA repair, excessive PARP1 activity is known to induce a form of cell death named parthanatos ^[23]^. The most widely accepted mechanism of parthanatos involves the translocation of the mitochondrial protein AIF to the nucleus triggered by PAR release, followed by AIF-mediated DNA degradation ^[24–26]^. This mechanism differs fundamentally from apoptosis, as it is strictly PARP1-dependent and caspase-independent, and there is evidence for cross-regulation between these pathways, as apoptosis involves caspase-dependent cleavage of PARP1 ^[27]^. Parthanatos has been implicated in several pathological processes, such as myocardial infarction, stroke, sepsis, and neurodegenerative diseases such as Parkinsons and Alzheimers diseases, and pharmacological inhibition or genetic deletion of PARP1 were shown to be cytoprotective in several model systems studied over the last decades ^[28–39]^. Since ADP-ribosylation consumes NAD^+^, the earliest proposed mechanisms of PARP-dependent cell death were centered around energetic deficiencies, particularly NAD^+^ and/or ATP depletion ^[27,40–43]^. Indeed, the depletion of NAD^+^ and of ATP were already observed when PARP1-dependent cell death was first described in the 1980s ^[40]^, and their occurrence in parthanatos-dependent pathological processes is well documented ^[32,33,35,37]^. However, the exact molecular mechanism by which hyperactivation of PARP1 leads to parthanatos and a clear sequence of events driving PARP1-dependent ATP depletion is currently lacking ^[44]^. The role of the hydrolases PARG and ARH3 in parthanatos is particularly unclear, with the literature pointing to conflicting effects of these enzymes on parthanatos execution ^[44–46]^.

In this work, we show that cell death by parthanatos is strictly dependent on PARG activity and is unaffected by ARH3 loss. While PARP1 hyperactivity is necessary and sufficient to drive NAD^+^ depletion, PARG inhibition uncoupled NAD^+^ depletion from ATP depletion, indicating that PAR hydrolysis is required for ATP depletion and that parthanatos does not necessarily correlate with NAD^+^ depletion. During these experiments, we uncovered evidence for a new PARG isoform of 53 kDa, which we propose to name PARG53.

## Results

### MNNG induces parthanatos

To study parthanatos, we first established conditions in which cells died effectively and specifically by parthanatos, which we define as acute DNA damage-induced cell death (within 24h of treatment) that can be prevented by PARP1/2 inhibition with olaparib. As reported for other cell lines ^[24,47–50]^, the alkylating agent N-methyl-N′ nitro-N-nitrosoguanidine (MNNG) strongly induced cell death by parthanatos in RPE1 cells, as determined using three different methods to measure cell death/viability (Fig. 1A-C). XTT assays, which reflect cell viability by measuring mitochondrial function, showed a major reduction in viability caused by MNNG treatment, which was partially prevented by the PARP1/2 inhibitor olaparib (Fig. 1A). However, this result is confounded by the continued replication of cells in the untreated condition, which increases overall mitochondrial activity in the control well, and the fact that mitochondrial dehydrogenase activity could be affected during parthanatos without a direct impact on cell death. Using flow cytometry and fluorescence microscopy-based assays, both relying on propidium iodide (PI) staining to detect cell death based on cell membrane rupturing, MNNG caused significant levels of cell death, with around 30 to 50% of PI-positive cells after 24h, which was completely prevented by the PARP1/2 inhibitor olaparib or by PARP1 deletion (Fig. 1B-C). To determine the kinetics of protein ADP-ribosylation and ADPr hydrolysis under these conditions, we determined total ADP-ribosylation (both MAR and PAR) via western blotting. As observed with many other PARP1-activating DNA damaging agents, MNNG induced a wave of ADP-ribosylation that peaked at the earliest collected time point (15 min), and its removal was virtually complete by 60 min (Sup. Fig. 1A-B). MNNG treatment was not associated with caspase-dependent cleavage of PARP1, indicating that cell death was non-apoptotic, as expected (Sup. Fig. 1C-D). Thus, we conclude that MNNG is an effective stimulus to induce canonical PARP1-dependent parthanatos in RPE1 cells.

**Figure 1:**
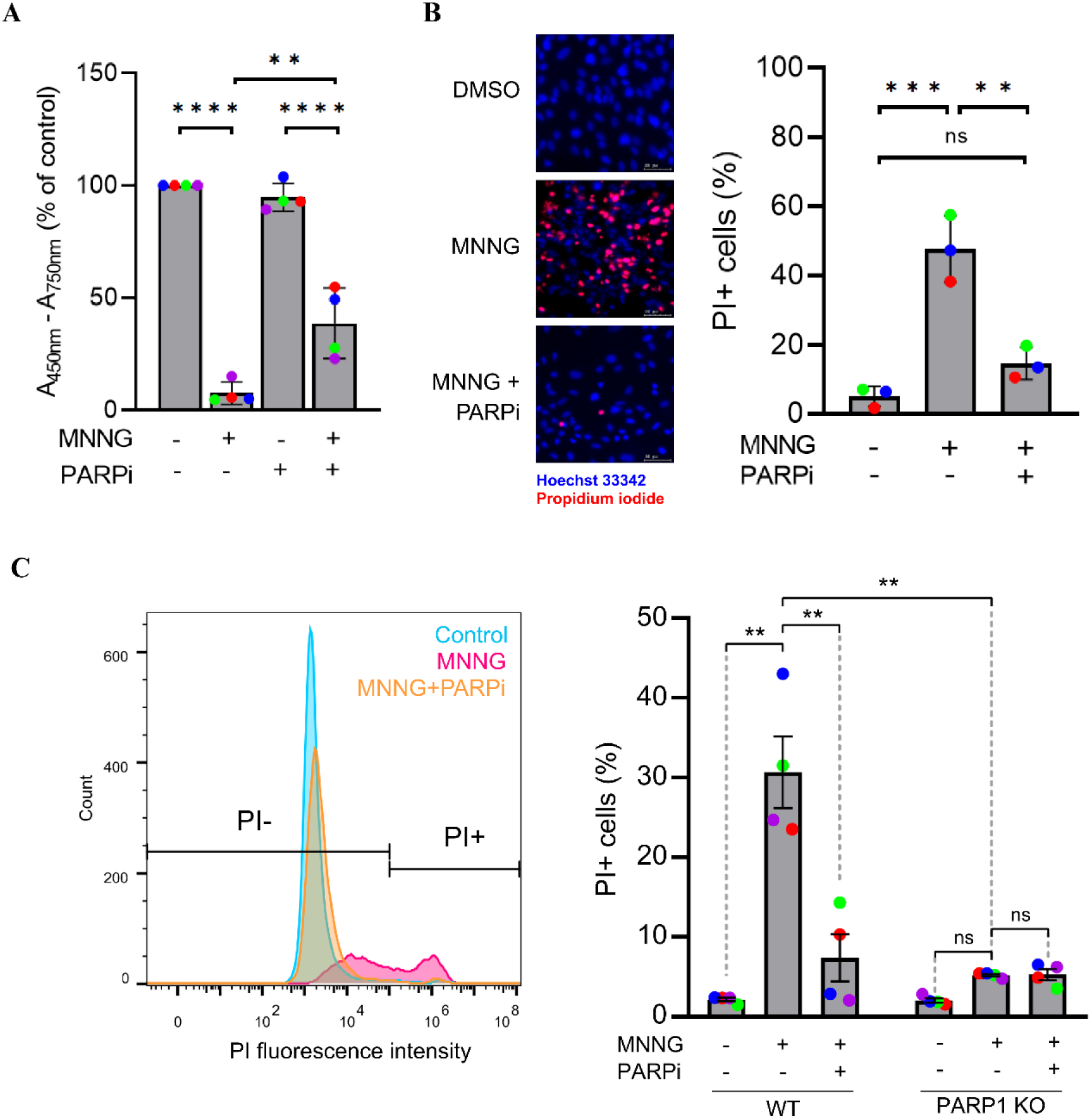
MNNG induces PARP1-dependent cell death in RPE1 cells. **A)** XTT cell viability assay in RPE1 cells treated for 24 hours with 250 µM MNNG and/or 10 µM olaparib (added one hour before and throughout the MNNG treatment), as indicated. **B)** Representative images (left) and quantification of PI-positive cells (right) using a fluorescence microscopy-based cell death assay. Cells were treated as in A (except that 5mM MNNG was used) and then incubated with Hoechst 33342 (blue) and propidium iodide (PI, red) for 10 min prior to image acquisition, without permeabilization or fixation. Scale bar = 10 µm. **C)** Representative histogram overlay (left) and quantification of PI+ cells (right) using a flow cytometry-based cell death assay. WT or PARP1 KO cells were treated as in A, and stained with PI without permeabilization or fixation. PI fluorescence intensity was quantified and cells above the shown threshold were considered PI-positive. All bar graphs show mean and SD of 3 or 4 independent experiments (coloured dots). Data were analysed using one-way ANOVA with Tukey’s test (ns = not significant, * = p < 0.05, ** = p < 0.005, *** = p < 0.0005, **** = p < 0.0001).

### Parthanatos requires PARG, but not ARH3

To study the contribution of PARG and ARH3 for parthanatos induction, we employed CRISPR/Cas9-generated knockout cells. While the ARH3 KO cells have been previously described ^[51]^, PARG KO cells were generated in this study, using an sgRNA targeting exon 3 of the *PARG* gene (Sup. Fig 2). In order to characterize the expected defects in ADP-ribosylation reversal in these knockout cell lines, we performed western blotting and immunofluorescence experiments detecting the kinetics of MAR/PAR formation and hydrolysis following MNNG treatment. Western blots showed a major defect in ADP-ribosylation reversal in PARG KO cells, with an increase in MAR/PAR compared to controls already evident at 15 min and a clear defect in ADPr hydrolysis by 60 min (Fig. 2A-B). A similar defect was also observed in cells treated with the PARG inhibitor PDD0017273 ^[52]^. As expected, ARH3 KO cells displayed a clear, but less pronounced, defect in ADPr hydrolysis, consistent with its known role in removing predominantly MAR ^[18,19]^. In these cells, clear accumulation of MAR/PAR signal over time was evident in the low molecular weight range, which likely represents histone MARylation (Fig. 2A-B). The treatment of ARH3 KO cells with PARG inhibitor led to the appearance of MAR/PAR signal even in untreated samples, indicating that endogenous ADPr is continuously hydrolysed by the concerted action of these enzymes. PARG inhibition also led to a reduction of the histone MAR/PAR signal observed in MNNG-treated ARH3 KO cells (Fig. 2A, lane 6 vs lane 12 - 15 kDa range), suggesting that, at least in response to MNNG treatment, most of the histone MARylation that accumulates in ARH3 KO cells requires the processing of PAR chains by PARG. To confirm this observation, we repeated the same experiment using recently developed reagents that specifically detect MAR ^[53,20]^. While MARylation was clearly induced in WT cells in the 15 kDa range after 15 min of MNNG treatment and then completely hydrolysed by 60 min, this signal was lost in both PARG KO or PARG inhibitor treated cells (Fig. 2C – top panel). In ARH3 KO cells, we again observed the accumulation of this 15 kDa MARylation signal over time, including the appearance of additional bands of higher molecular weight 60 min after MNNG treatment (Fig. 2C- top panel). When ARH3 KO cells were additionally treated with PARG inhibitor, this MARylation signal was again lost (Fig. 2C- top panel). Similar results were obtained using a site-specific antibody that detects histone H3 serine 10 MARylation, with ARH3 KO cells accumulating H3S10-MAR over time in a PARG-dependent manner. However, some residual H3S10 MARylation still accumulated in ARH3 KO cells treated with PARGi (Fig. 2C – bottom panel), indicating low levels of direct MARylation, in agreement with previous observations ^[20]^. To further characterize the defects in ADPr hydrolysis in response to MNNG in these cell lines, we performed a similar set of experiments detecting MAR/PAR via immunofluorescence. In control cells, MNNG induced nuclear MAR/PAR formation by 15 min, which was completely reversed by 60 min (Fig. 2D-E). Similar to the immunoblot results, loss of either ARH3 activity or PARG activity delayed ADPr hydrolysis, as expected (Fig. 2D-E). Interestingly, little to no difference in signal intensity was noticeable between PARG KO, ARH3 KO or PARG inhibitor treated cells in these experiments, likely due to the difficulty in distinguishing MAR from PAR in this experiment, due to the use of a pan-ADPr specific reagent (Fig. 2D-E). Collectively, these data indicate that MNNG treatment induces a wave of ADP-ribosylation that is rapidly hydrolysed by the combined activities of ARH3 and PARG, and confirm the functional defect in ARH3 and PARG activities in our knockout cell lines.

**Figure 2:**
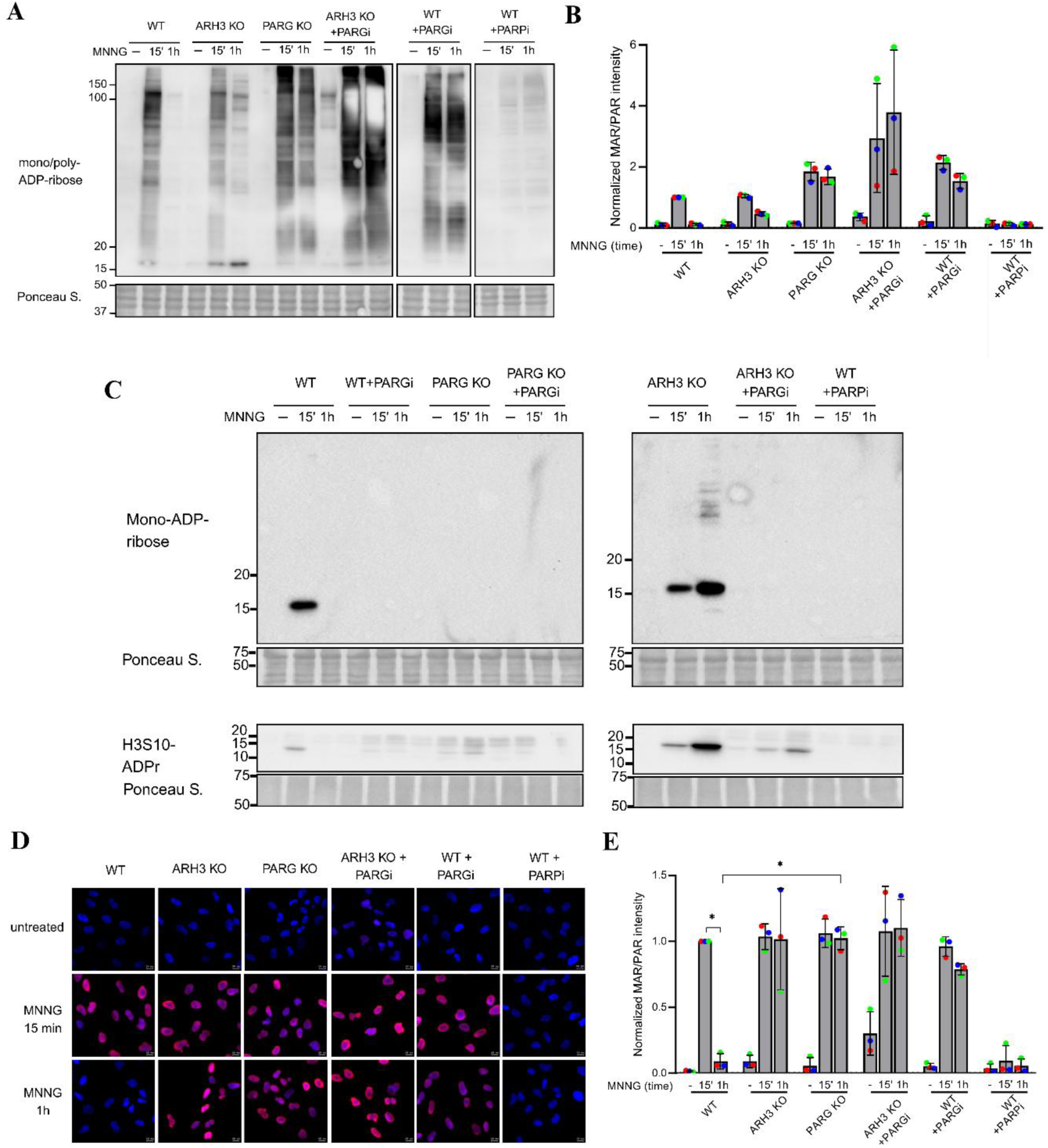
ARH3 and PARG are required for ADPr hydrolysis after MNNG treatment. **A)** Representative western blot for mono/poly-ADP-ribosylation (MABE1016) in RPE1 cells of the indicated genotypes treated or not with 250 µM MNNG for either 15 min or one hour. 10 µM olaparib (PARPi) or 10 µM PDD0017273 (PARGi) were added one hour before and during the MNNG treatments. **B)** Quantification of MAR/PAR signal from immunoblots performed as in A, normalized to Ponceau S signal, and then normalized to the WT 15’ sample for each replicate, which was assigned an arbitrary value of 1. **C)** Western blot for mono-ADP-ribosylation (AbD43647 - top) and Histone H3 serine 10-mono-ADPr (H3S10-ADPr) (AbD33644 - bottom) in cells treated as in A. **D)** Representative immunofluorescence microscopy images of cells treated as in A, stained with DAPI (blue) and mono/poly-ADPr (MABE1016 - red). Scale bar = 10 µm. **E)** Quantification of the mean MAR/PAR fluorescence intensity per nucleus from replicate experiments performed as in D. Thousands of nuclei were quantified per replicate, and the mean signal per sample was normalized to the respective WT 15’ sample, which was assigned an arbitrary value of 1. **B and E)** Graphs depict mean and SD of three independent experiments (coloured dots). Data were analysed using one-way ANOVA with Tukey’s test (* = p < 0.05).

Next, we used this isogenic set of cell lines to investigate the contribution of ARH3 and PARG towards cell death by parthanatos, measuring MNNG-induced cell death by flow cytometry. Similar to Fig. 1C, close to 50% of WT RPE1 cells became PI-positive 24 h after MNNG treatment, and this cell death was completely prevented by PARP1/2 inhibition with olaparib (Fig. 3A-B and Sup. Fig. 3A), indicating that these cells are dying by parthanatos under these conditions. ARH3 KO had no effect on cell death, suggesting that ARH3 is not required for parthanatos execution in this system. In contrast, PARG inhibition completely prevented cell death in these experiments, indicating that PARG activity is required for parthanatos(Fig. 3A-B). To confirm this effect in other cell lines, we induced parthanatos by high-dose MNNG treatment of Hela and A549 cells, and again observed that PARG inhibition prevented parthanatos execution (Sup. Fig. 3C-D). Taken together, these results suggest that while ARH3 is dispensable for cell death by parthanatos, complete inhibition of PARG activity prevents cell death by this mechanism, implying that both the formation of PAR chains by PARP1 and their hydrolysis by PARG are required to trigger parthanatos.

**Figure 3:**
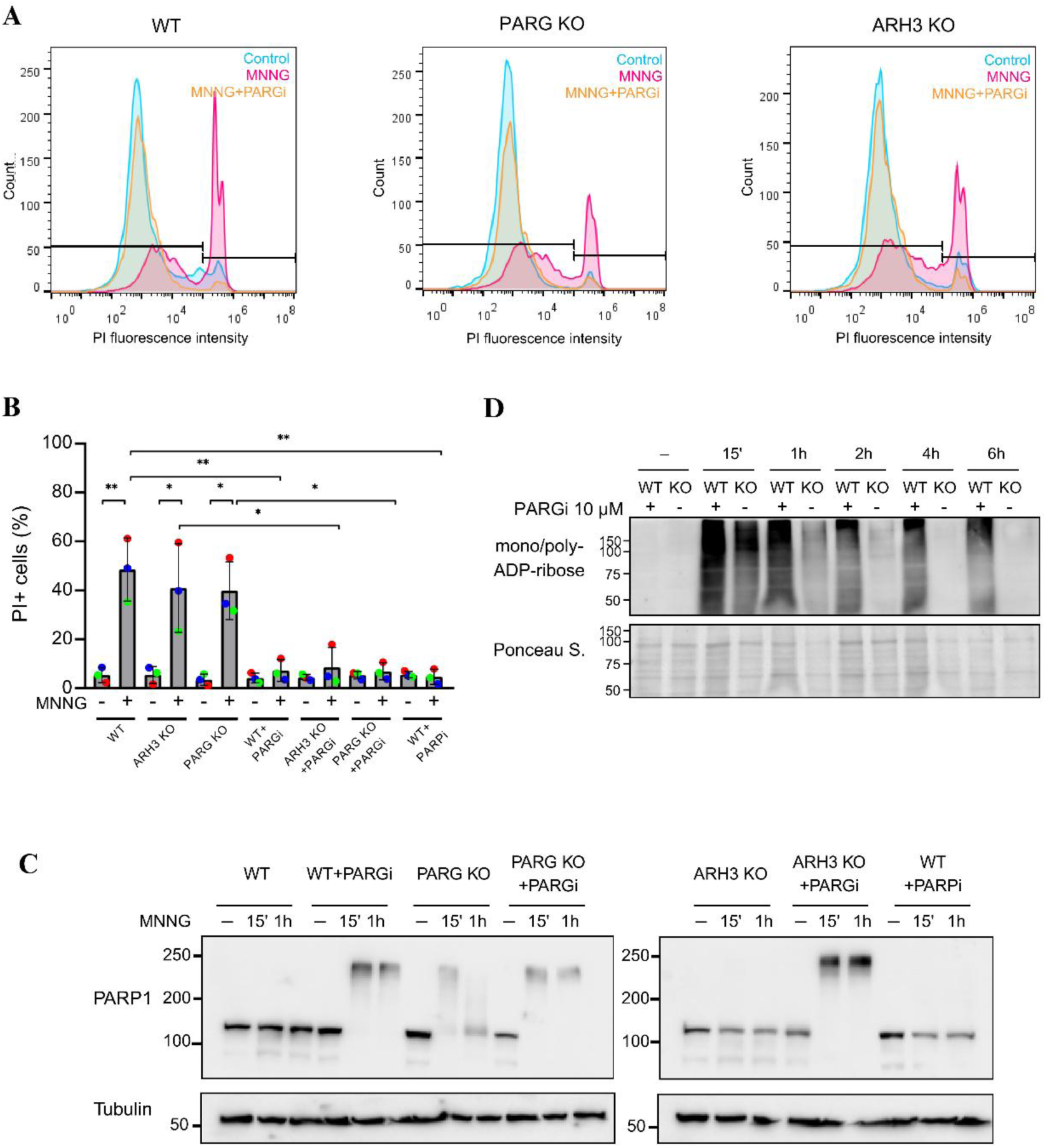
PARG, but not ARH3, is necessary for cell death by parthanatos. **A)** Representative histogram overlay and **B)** quantification of PI-positive cells using flow cytometry-based cell death assay. RPE1 cells of the indicated genotypes were treated or not with 250 μM MNNG for 24h and 10 µM olaparib (PARPi) or 10 µM PDD0017273 (PARGi), which were added one hour before and during the MNNG treatments. Graphs depict mean and SD of three independent experiments (coloured dots). Data were analysed using one-way ANOVA with Tukey’s test (* = p<0.05; ** = p<0.01). **C)** Representative western blot for PARP1 protein in RPE1 cells of the indicated genotypes treated or not with 250 µM MNNG for either 15 min or one hour. 10 µM olaparib (PARPi) or 10 µM PDD0017273 (PARGi) were added one hour before and during the MNNG treatments. **D)** Representative western blot for mono/poly-ADP-ribosylation (MABE1016) in WT RPE1 cells treated continuously with 10 µM PDD0017273 (PARGi) or in PARG KO cells, after treatment with 250 μM MNNG for the indicated times.

In the above experiments, we were surprised to find that our PARG KO cells were still dying by parthanatos, which differed from the effect observed with PARG inhitor. Indeed, the effect of PARG inhibitor was still observed in PARG KO cells (Fig. 3A-B), indicating either that there is residual PARG activity in our PARG knockout cells, or that the PARG inhibitor was engaging additional cellular targets beyond PARG to mediate this effect. Therefore, we sought to more stringently test the levels of PARG activity in our KO cells. Given that PARP1 activation leads to its auto-PARylation, and that the size of the PAR chain attached to PARP1 would be reflected on its electrophoretic mobility, we reasoned that immunoblots probing for PARP1 protein (rather than MAR/PAR) could serve this purpose. Indeed, whereas MNNG treatment did not substantially impact PARP1 migration in control cells at any time point (despite detectable MAR/PAR formation at 15 min - see Fig. 2A), treatment with PARG inhibitor led to a striking MNNG-induced shift of the whole cellular complement of PARP1 protein by over 100 kDa, indicating the accumulation of hyperPARylated PARP1 in these cells (Fig 3C). In our PARG KO cells, we observed a similar mobility shift of PARP1 15 min after MNNG treatment, confirming that these cells are defective in PARG activity compared to controls. However, there was a clear return of a substantial portion of PARP1 molecules to their initial electrophoretic mobility over time, indicating active removal of PAR chains in the PARG KO cells, albeit at a slow rate. This effect was again inhibited when these cells were treated with PARG inhibitor, indicating that this reflects low levels of PARG activity in PARG KO cells (Fig. 3C). To confirm this result, we monitored MAR/PAR levels for up to six hours after MNNG treatment in PARG KO cells and compared this to WT cells treated with PARG inhibitor. While PARG inhibition prevented ADPr hydrolysis over the whole period, there was a clear reduction in MAR/PAR levels in our PARG KO cells over time (Fig. 3D). These data indicate that our PARG KO cell line is a hypomorph, and that residual PARG activity in these cells explains the discrepancy between parthanatos induction in these cells compared to PARG inhibitor treated cells.

### Low levels of PARG activity are sufficient for parthanatos

The above data indicated that the small amount of residual PARG activity still present in our PARG KO cells was sufficient to induce parthanatos. To corroborate this finding, we treated cells with decreasing concentrations of PARG inhibitor and probed for residual MAR/PAR 1h and 4h after MNNG treatment (Fig. 4A-B). As expected, PARG KO cells retained substantial amounts of MAR/PAR 1h after MNNG treatment, whereas MAR/PAR was completely hydrolysed in WT cells by this time point (Fig. 4A-B and Sup. Fig 4). Similar to Fig. 3D, continued PAR hydrolysis in the PARG KO cells led to complete removal of the MAR/PAR signal by 4h. In contrast, treatment of WT cells with PARG inhibitor at 10 µM (the concentration used for all experiments so far) largely prevented PAR hydrolysis up to 4h after treatment. Although concentrations of PARG inhibitor in the 1-3 μM range still reduced PAR degradation compared to controls at 1h (Sup. Fig. 4), PAR hydrolysis was incompletely inhibited in these cells, as the MAR/PAR signal continued to decline over time (Fig. 4A-B). To corroborate these findings, we determined the effect of these doses of PARG inhibitor on MNNG-induced PARP1 hyperPARylation (similar to Fig. 3C). Again we observed that PARG KO cells had a clear defect in the hydrolysis of PARP1 autoPARylation compared to controls, but hydrolysis was still observed over time (Fig. 4C-D). While 10 µM PARG inhibitor completely prevented PAR hydrolysis in this assay, lower doses in the 1-3 µM range were insufficient to completely block PARG activity (Fig. 4C-D). Next, we determined cell death by parthanatos under these conditions using flow cytometry. Similar to Fig. 2, MNNG induced cell death in both WT and PARG KO cells, which was prevented by PARG inhibitor at 10 µM (Fig. 4E-F). However, when PARG inhibitor was used in the 1-3 µM range, cell death by parthanatos was still observed (Fig. 4E-F). These results indicate that low levels of PARG activity are sufficient to trigger parthanatos, thus providing an explanation for why our hypomorphic PARG knockout cell line is proficient for parthanatos.

**Figure 4:**
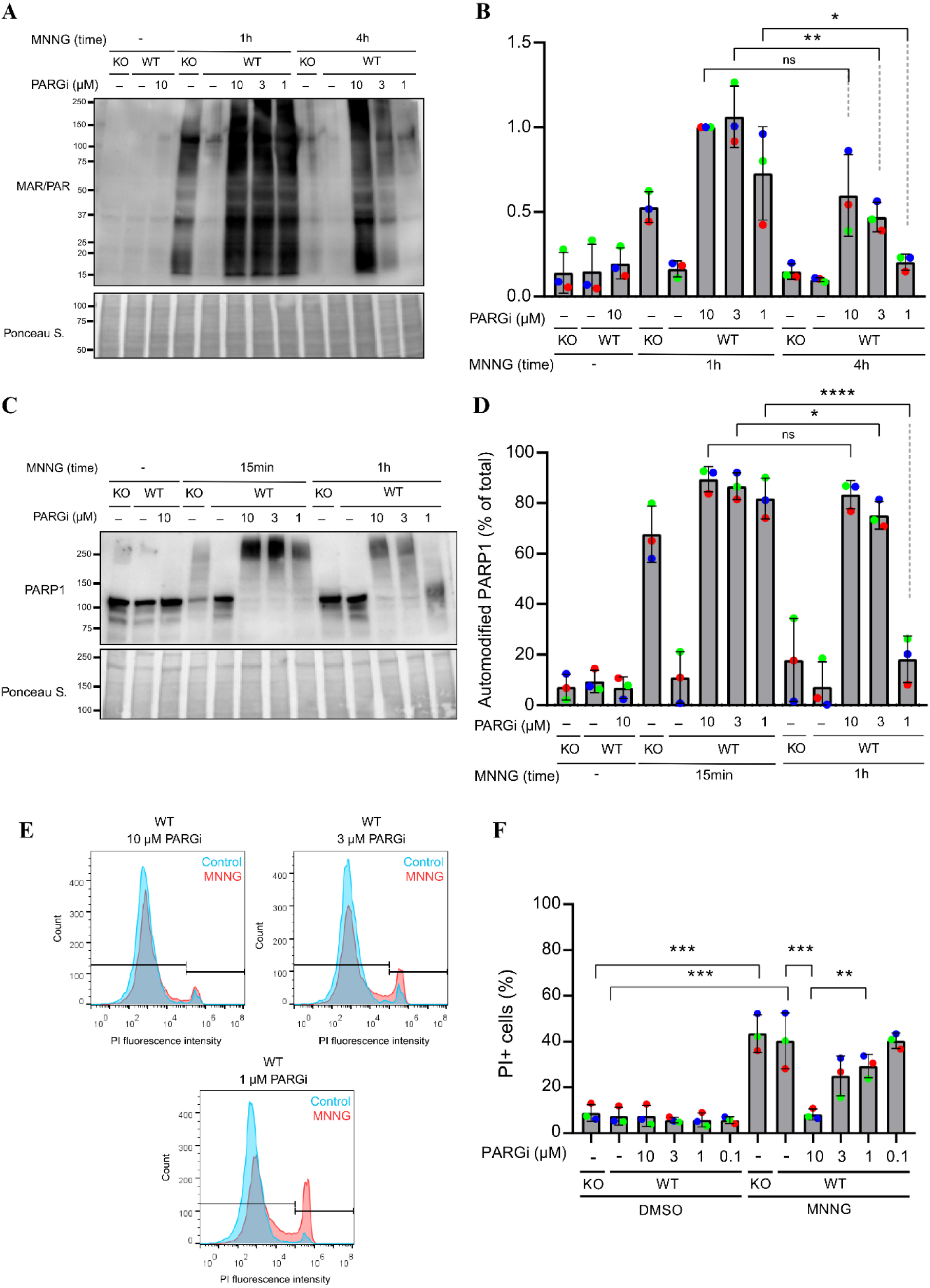
Low levels of PARG activity are sufficient for cell death by parthanatos. **A)** Representative western blot and **(B)** quantification of mono/poly-ADP-ribosylation (MABE1016) in PARG KO cells or WT RPE1 cells treated with the indicated concentrations of PDD0017273 (PARGi) and DMSO control (-) or 250 μM MNNG for one hour (1h) or four hours (4h). **C)** Representative western blot for PARP1 protein and **(D)** quantification of hypermodified PARP1 in PARG KO cells or WT RPE1 cells treated with the indicated concentrations of PDD0017273 (PARGi) and with DMSO control or 250 μM MNNG for 15 minutes (15min) or one hour (1h). **(E)** Representative histogram overlays and **(F)** quantification of PI-positive cells using flow cytometry-based cell death assay. WT RPE1 cells or PARG KO cells were treated with 250 μM MNNG for 24 hours. PDD0017273 (PARGi) was added one hour before and during the MNNG treatments, as indicated. Graphs depict mean and SD of three independent experiments (coloured dots). Data were analysed using two-way ANOVA with Tukey’s test (** = p < 0.005, *** = p < 0.0005).

### A new PARG isoform

The next question we sought to address was the origin of the residual PARG activity in our PARG KO cell line. These cells were generated using an sgRNA targeting exon 3, and were derived by clonal expansion. PARG protein levels were undetectable in the KO cells at the expected ∼100 kDa range, and further siRNA depletion of PARG did not impact any of the remaining bands detected by our PARG antibody (Fig. 5A). Sanger sequencing confirmed a homozygous 4 nt deletion at the Cas9 cleavage site, indicating successful editing of the locus (Sup. Fig. 2). During the course of this work, the Junjie Chen group showed that a full PARG KO is not viable, unless cells are grown in the presence of PARP1/2 inhibitor, indicating that PARG is an essential gene ^[54]^. Similar to our observations, they found that PARG KO clones generated using sgRNAs targeting exon 3 (in the vicinity of our sgRNA) or targeting exon 7 were viable due to residual PARG activity, while targeting of later exons (from exon 9 onwards) was necessary to completely prevent PARG activity ^[54]^. Given that the main PARG isoforms encoding PARG111, PARG102 and PARG99 are all identical from exon 3 onwards and that even the annotated splice isoforms encoding putative PARG60 and PARG55, which are not thought to be catalytically active, are identical from exon 6 onwards (Fig. 5B), none of the annotated PARG isoforms could explain the residual PARG activity both of our groups identified.

**Figure 5:**
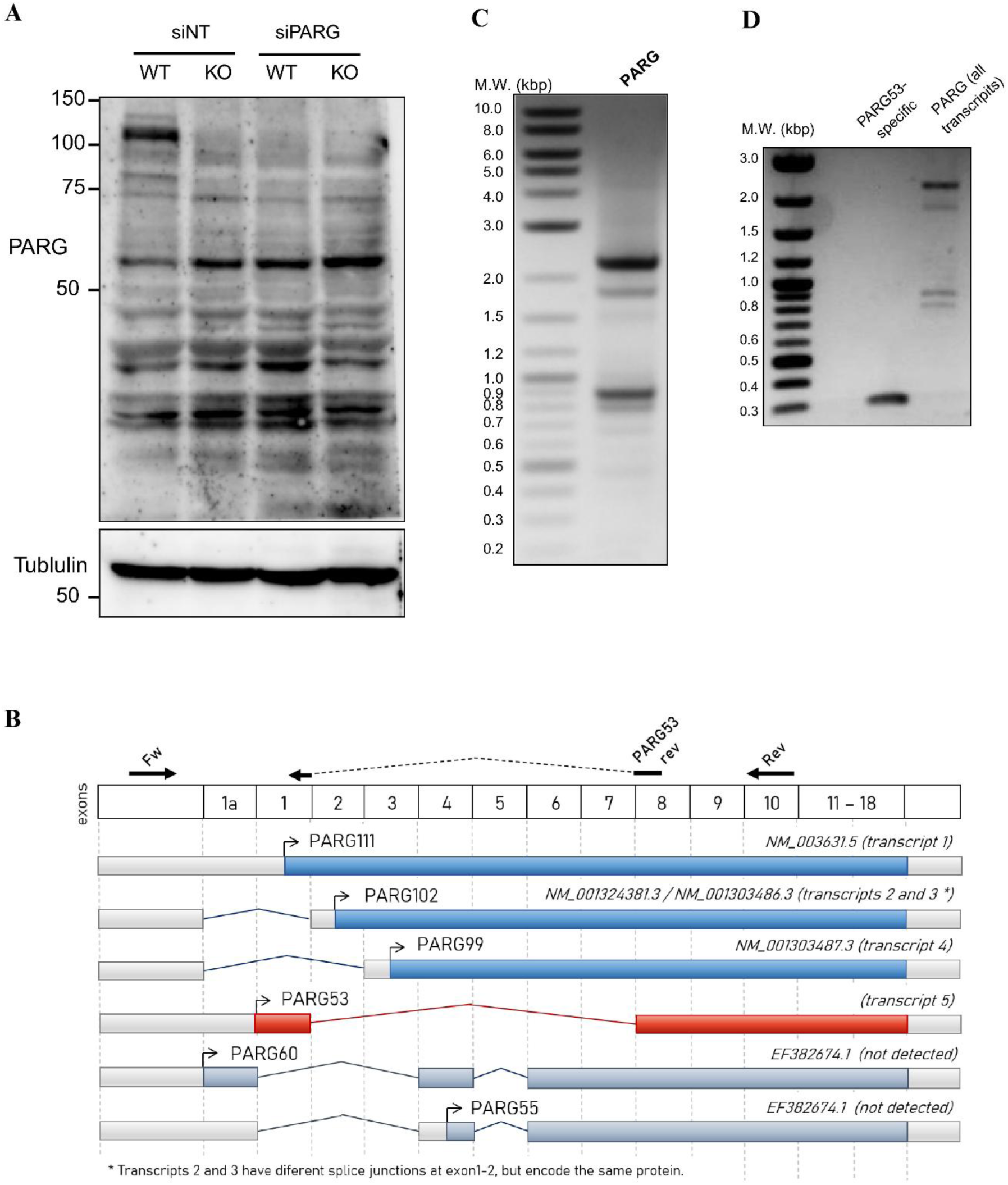
PARG53 is a new PARG isoform. **A)** Representative western blot for PARG protein in WT RPE1 cells or PARG KO cells transfected either with non-targeting control (siNT) or a pool of siRNAs targeting PARG (siPARG). **B)** Schematic representation of transcripts encoding known and putative PARG isoforms. Exons 1 to 18 are shown in white boxes across the top (not to scale), each transcript is shown as a grey bar, labelled with the respective annotated transcript and the transcript numbering from Table1. Encoded ORFs are coloured blue for canonical PARG isoforms, red for the new isoform identified here, and light blue for isoforms described in the literature that we were unable to detect. Locations of the PCR primers used for panels C and D are shown across the top (not to scale) **C)** Representative image of an agarose gel showing the products of a PCR reaction using RPE1 cDNA as template and primers targeting the 5’UTR (Fw) and exon 10 (Rev) of *PARG* **D)** Similar to C, agarose gel showing the products of PCR reactions using the same 5’UTR forward primer combined either with a PARG53-specific reverse primer spanning the exon 1-8 junction (left - expected size 297bp) or the exon 10 reverse primer (right).

To explore the full spectrum of splice variants within the 5′ end of PARG transcripts, we performed PCR on cDNA obtained from WT RPE1 cells using a forward primer targeting the 5′-UTR and a reverse primer within exon 10 (Fig. 5B). Several PCR products of various sizes were visible on an agarose gel, up to the 2.2 kb size expected for the reference transcript encoding PARG111, confirming extensive alternative splicing of PARG transcripts (Fig. 5C). This population of PCR products was then subjected to Oxford Nanopore sequencing, which is a long-read sequencing technology, and therefore allows more reliable detection of splice variants. Reads containing similar splicing patterns were assembled into a total of 16 different putative transcripts, whose overall sizes and relative abundance matched the pattern observed in the agarose gel, and included sequences that matched annotated *PARG* gene transcripts (Table 1 and Fig. 5C). The longest ORFs predicted to be encoded by each of these transcripts are also shown in Table 1, and included transcripts for PARG111, 102 and 99, as expected. However, many of these assembled transcripts did not align to the *PARG* gene, but instead most closely matched the sequence of the *PARGP1* pseudogene. This pseudogene contains a large portion of the 5′ end of the *PARG* gene, including both of the primer sequences we used for PCR amplification, but is not predicted to encode a functional protein, as it does not contain exons 13 to 18, which encode a large portion of the PARG catalytic domain. While we could not find any evidence for transcripts encoding full length PARG60 or PARG55, several transcripts that aligned to the pseudogene harboured the exon 1/1a-4 junction, followed by an exon 4-6 junction, which is the splicing pattern that characterizes the transcripts for these putative isoforms (Fig. 5B). A more refined analysis of this sequencing data, based on the identification of diagnostic polymorphisms that distinguish the genomic origin of each individual read, revealed that none of the reads from the *PARG* gene contained this e1-e4-e6 splicing pattern, and a large portion of reads originating from the pseudogene did (Sup. Fig. 5 and Supplementary Information). This data indicates that isoforms PARG60 and PARG55 are likely to have been originally identified based on PCR amplification from the *PARGP1* pseudogene, rather than from the true *PARG* gene.

**Table 1:**
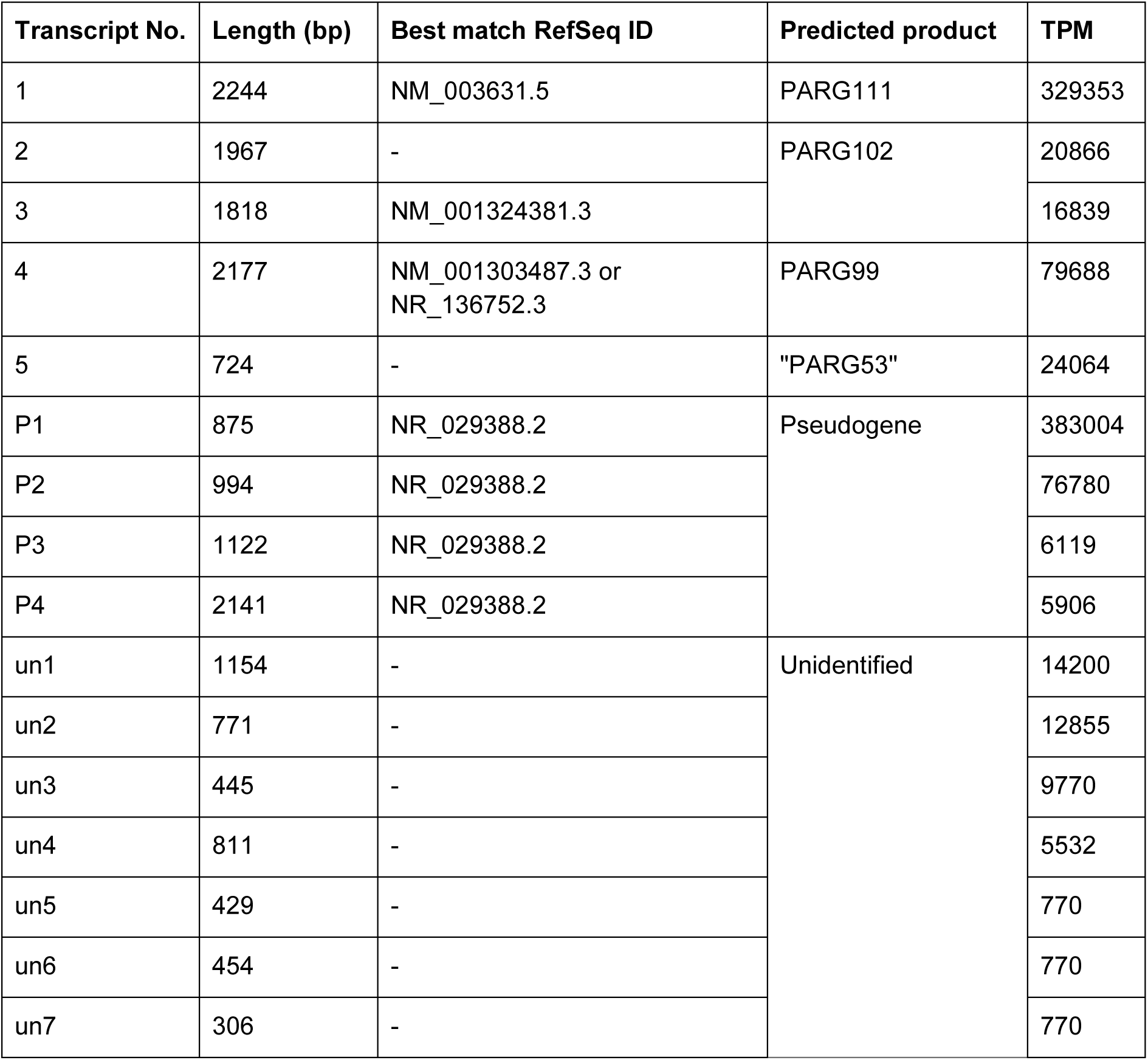
List of assembled transcripts from the long-read sequencing analysis of PCR products shown in Fig. 5C. Raw reads were assembled into the 16 shown transcripts using StringTie. “Best match RefSeq IDs” refer to known transcripts with 100% identity, largest e-value, and largest query-cover determined by nucleotide BLAST. “Predicted products” are the largest ORF for each assembled transcript. Length (in bp) and Transcripts per million (TPM) values are shown for each individual transcript.

| Transcript No. | Length (bp) | Best match RefSeq ID | Predicted product | TPM |
| --- | --- | --- | --- | --- |
| 1 | 2244 | NM_003631.5 | PARG111 | 329353 |
| 2 | 1967 | - | PARG102 | 20866 |
| 3 | 1818 | NM_001324381.3 |  | 16839 |
| 4 | 2177 | NM_001303487.3 or<br>NR_136752.3 | PARG99 | 79688 |
| 5 | 724 | - | "PARG53" | 24064 |
| P1 | 875 | NR_029388.2 | Pseudogene | 383004 |
| P2 | 994 | NR_029388.2 |  | 76780 |
| P3 | 1122 | NR_029388.2 |  | 6119 |
| P4 | 2141 | NR_029388.2 |  | 5906 |
| un1 | 1154 | - | Unidentified | 14200 |
| un2 | 771 | - |  | 12855 |
| un3 | 445 | - |  | 9770 |
| un4 | 811 | - |  | 5532 |
| un5 | 429 | - |  | 770 |
| un6 | 454 | - |  | 770 |
| un7 | 306 | - |  | 770 |

Our assembly also revealed a transcript which contains a novel exon 1 - exon 8 junction that is not currently found in any database and is predicted to encode a 53 kDa protein, which we propose to call PARG53. We estimated the abundance of this transcript to be at roughly the same level as transcripts encoding PARG102 and PARG99 (Sup. Fig. 5 and Supplementary Information), and have identified several raw reads that originated from the *PARG* gene and contain the novel splice junction (Sup. Fig. 5 and Supplementary Information). To further investigate the existence of this isoform, we performed PCR on RPE1 cDNA using a primer targeting the novel exon 1-8 junction. This resulted in an amplicon of the expected size (297bp) (Fig. 5D), providing additional evidence for the existence of this isoform. Finally, we sought to specifically knockdown PARG53 using custom siRNAs targeting this exon boundary. However, our siRNAs did not result in the knockdown of any bands near the 50 kDa region, nor were they specific for these shorter isoforms (Sup. Fig. 6). While the full characterization of this novel isoform is beyond the scope of this work, its identification provides an explanation for the residual PARG activity in our PARG KO cells.

### PARP1-induced ATP depletion (but not NAD^+^ depletion) is dependent on PARG activity

Our results so far show that low levels of PARG activity are necessary for parthanatos execution, but do not address how exactly PARG activity contributes to this pathway. Therefore, we investigated the impacts of impaired PARG activity on the energetic collapse associated with parthanatos. For this, we first determined the activation of the AMP-activated kinase (AMPK), which responds to changes in AMP/ATP ratios, by western blotting ^[55]^. Treatment with MNNG caused an increase in AMPK activation, as measured by phosphorylation on Thr172, within 1h (Fig. 6A and Sup. Fig. 7A). This AMPK phosphorylation was prevented by treatment with olaparib or PARP1 deletion, confirming its dependency on PARP1 hyperactivation (Fig. 6A and Sup. Fig. 7B). ARH3 KO or PARG KO had no effect on AMPK activation, whereas PARG inhibitor treatment completely prevented MNNG-induced AMPK-Thr172 phosphorylation (Fig. 6A). To confirm that this discrepancy between PARG inhibitor and PARG KO cells was again due to low levels of residual PARG activity, we used decreasing doses of PARG inhibitor, and observed activation of AMPK at PARG inhibitor concentrations in the 1-3 µM range (Sup. Fig. 7C). Thus, PARP hyperactivation induces AMPK activation and, in a manner analogous to cell death, at least a low level of PARG activity is necessary for such effect.

**Figure 6:**
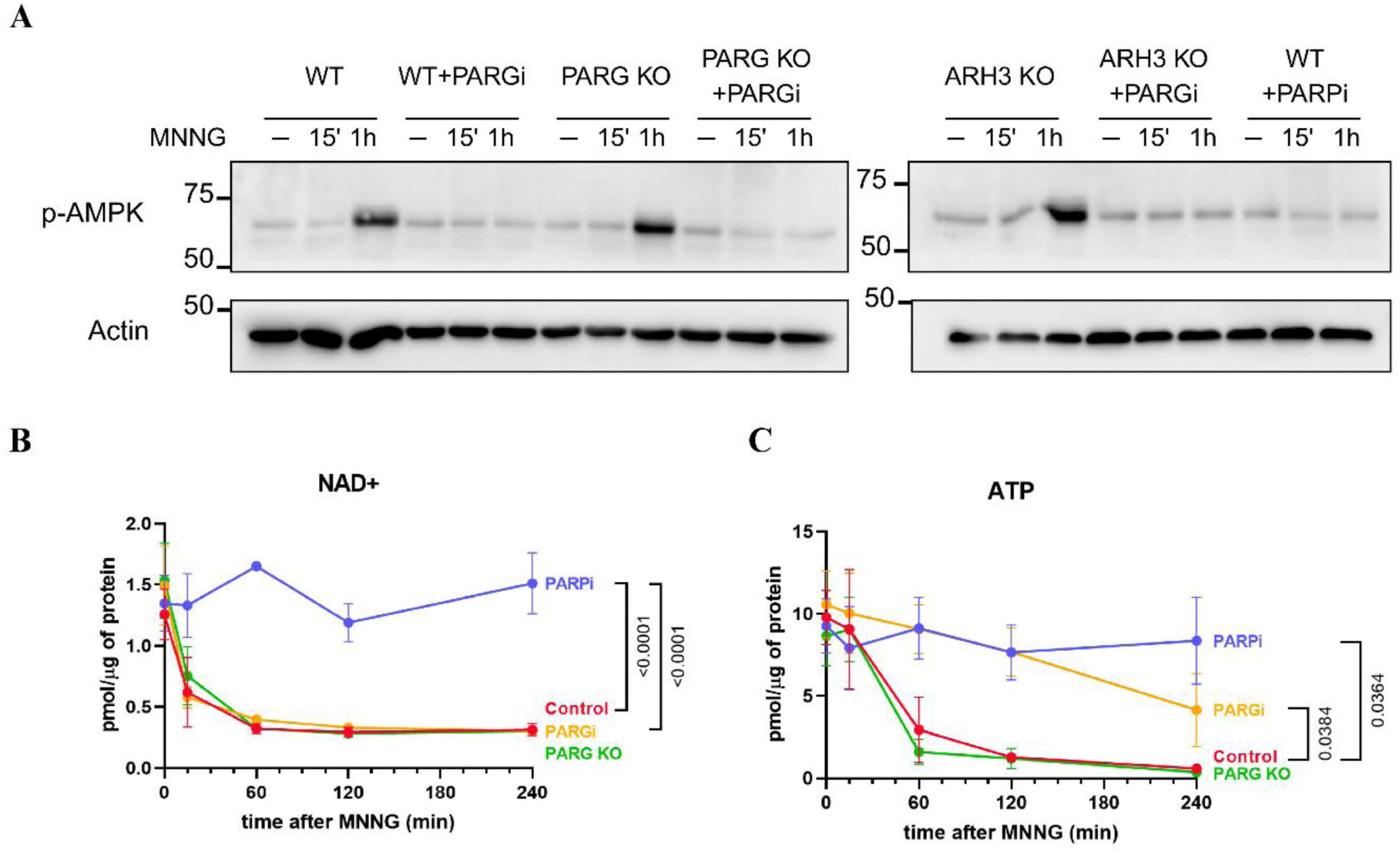
PARG activity is required for ATP depletion during parthanatos. **A)** Representative western blot for AMPK phosphorylation at threonine 172 and actin loading control in RPE1 cells of the indicated genotypes treated or not with 250 µM MNNG for either 15 min or one hour. 10 µM olaparib (PARPi) or 10 µM PDD0017273 (PARGi) were added one hour before and during the MNNG treatments. **B-C)** Quantification of NAD^+^ (B) and ATP (C) by HPLC-UV in RPE1 WT or PARG KO cells treated with DMSO control (‘0 min’) or treated with 250 µM MNNG for 15, 60, 120 or 240 min. 10 µM olaparib (PARPi) or 10 µM PDD0017273 (PARGi) were added to WT cells one hour before and during the MNNG treatments, as indicated. Graphs show mean and SEM of 4 independent experiments. For statistical analysis, the area under the curve for each condition was compared using one-way ANOVA and Tukey’s test. p-values are indicated for each comparison.

To more directly quantify the levels of ATP, ADP and AMP during MNNG treatment, as well as the NAD^+^ and NADH pools, we employed a HPLC-based method we recently established ^[56]^. As expected, MNNG treatment induced a profound and sustained depletion of both NAD^+^ and ATP, which were both prevented by olaparib treatment, indicating that they are PARP1-dependent (Fig. 6B-C). Notably, whereas NAD^+^ depletion was already evident by 15 min (Fig. 6B), ATP depletion was only observed after 1h (Fig. 6C), in agreement with our western blotting data showing PAR formation by 15 min (Fig. 2A) and AMPK activation at 1h (Fig. 6A). While these effects still occurred in PARG KO cells, PARG inhibition largely prevented ATP depletion, whereas NAD^+^ depletion was unaffected (Fig. 6B-C). While ADP and NADH levels remained fairly constant throughout this experiment (Sup. Fig. 8A-B), an apparent increase in AMP was observed after 1h, which was also PARP and PARG dependent (Sup. Fig. 8C), but did not reach statistical significance. Taken together, these results suggest that PARG activity is necessary for the depletion of cellular ATP pools caused by PARP1 hyperactivation - possibly accompanied by a rise in AMP - while PARP1 activity alone is sufficient to deplete cellular NAD^+^. Therefore, ATP depletion can be uncoupled from NAD^+^ depletion, indicating that one is not a direct consequence of the other.

## Discussion

Parthanatos is a form of cell death that results from DNA damage-induced hyperactivation of PARP1, and is characterized by this strict PARP1-dependency. However, the contribution of the main PAR hydrolases ARH3 and PARG for this type of cell death is less well established ^[44]^. The data presented here indicate that while ARH3 is largely dispensable for parthanatos execution, PARG activity is necessary for cell death by this pathway. The results concerning ARH3 differ from previous studies in which ARH3 deletion was shown to sensitize to cell death by parthanatos, including in a mouse model of ischaemia-reperfusion injury ^[45,46]^. While we cannot rule out a minor contribution of ARH3, we focused on exploring the molecular mechanisms underlying the strong effect observed with PARG inhibition, which completely suppressed MNNG-induced cell death by parthanatos in different human cell lines (Fig. 3 and Sup. Fig 3), as there is conflicting evidence in the literature regarding the contribution of PARG for parthanatos ^[44]^.

Due to a surprising phenotypic difference between our PARG KO cells and PARG inhibitor treatment, we uncovered that although PARG activity is strictly necessary for parthanatos execution, low levels of PARG function are sufficient for cell death induction (Fig. 3). This was observed both in our PARG KO cells, which have low levels of residual PARG activity (Fig. 3), or by reducing the concentration of PARG inhibitor to slightly lower levels (Fig. 4). This requirement for full PARG inhibition may have contributed to the conflicting evidence in the literature regarding the role of PARG for parthanatos ^[44]^, and indicates that somewhat slower kinetics of PAR hydrolysis are sufficient for cell death by parthanatos to be induced. Given that PARG is required for hydrolysis of PAR chains from autoPARylated PARP1 (Fig. 3 and ^[57]^), and that automodified PARP1 has reduced affinity for DNA breaks ^[58]^, one possible explanation for this effect is that full PARG inhibition leads to the accumulation of essentially inactive hypermodified PARP1, whereas residual PARG activity is sufficient to sustain a high enough level of PARP1 activity for parthanatos induction. Although we cannot rule out this possibility, complete PARG inhibition did not prevent the PARP1-dependent depletion of NAD^+^ pools (Fig. 6), suggesting that the consumption of NAD^+^ by PARP1 was normal under these conditions.

PARG activity is also essential for formation of the products of PAR hydrolysis, either in the form of free (i.e. not protein-linked) PAR chains produced by PARG endoglycohydrolase activity or ADP-ribose monomers generated by PARG exoglycohydrolase. Indeed, several reports indicate that free PAR chains are a crucial mediator of parthanatos ^[25,26,59]^, which could in principle explain the parthanatos defect after complete PARG inhibition. However, it was recently demonstrated that PARP1 can directly generate free PAR chains by catalysing PAR formation upon an NAD^+^ substrate molecule, instead of a protein ^[60]^. While PARG inhibition should prevent the formation of free PAR chains originating from protein-linked PAR, it is not expected to impact the formation of these NAD^+^-linked PAR chains, which were actually increased upon PARG inhibitor treatment ^[60]^. Our results would therefore indicate that NAD^+^-linked PAR chains are unlikely to mediate cell death by parthanatos, but further work will be required to test this idea. Another possibility that warrants further exploration is the role of PARG-dependent ADP-ribose monomers in mediating cell death, given that PARG is thought to preferentially act as an exoglycohydrolase ^[16,61]^.

In this context, it is important to point out that free PAR chains are thought to promote the release of AIF from the mitochondria to the nucleus, which is generally viewed as a key event in parthanatos induction ^[23,24,26,59]^. While we did not observe AIF translocation during parthanatos in RPE1 cells, in agreement with previous reports ^[62]^, we identified several interventions that prevent PARP1-dependent AIF translocation in other cell types, including PARG inhibitor treatment, and will report on these results elsewhere (manuscript in preparation).

The abovementioned difference between our PARG KO cells and PARG inhibitor treated cells led us to identify a novel PARG isoform predicted to have 53 kDa, which we propose to name PARG53. This isoform is encoded by a novel transcript in which exon 1 is spliced into exon 8, skipping exons 2 to 7 entirely (Figure 5 and Supplementary Information). Although we were unable to definitively detect this protein by western blotting (Fig. 5 and Sup. Fig. 3), evidence for a PARG isoform in this size range has been reported in the literature in the past, and was ascribed to two different alternatively spliced transcripts encoding PARG60 and PARG55 ^[11,12]^. The discovery of human PARG60/55 was motivated at the time by observations that while PARG knockout was embryonic lethal in mice, a hypomorphic allele in which exons 2 and 3 were deleted (which did not impact expression of the 50-60 kDa isoform) was compatible with life ^[47,63]^. With the advent of CRISPR/Cas9 technology, a similar effect was observed with the human *PARG* gene, namely that sgRNAs targeting exons 3 to 7 led to clones with residual PARG activity, while full PARG knockout cells generated with sgRNAs targeting later exons were not viable (this work and ^[54]^). Our discovery of PARG53, combined with the observation that PARG55 and PARG60 are likely to have been derived from transcripts originating from the *PARGP1* pseudogene, reconciles these findings and indicates that PARG53 is the small PARG isoform responsible for residual PARG activity in hypomorphic mutants in which the 5’end of the *PARG* gene is targeted.

The clear effects of the PARG inhibitor PDD0017273 in preventing parthanatos in our PARG KO cells ^[52]^ could be explained by off-target effects of the inhibitor towards an unknown alternative PAR hydrolase. However, given that our PARG KO clone clearly has residual PAR hydrolase activity (Fig. 3), that full PARG KO cells should not be viable ^[54]^, and that we identified a new PARG isoform that could provide residual PARG activity in our KO clones (Fig. 5), a more parsimonious interpretation of our data is that the PARG inhibitor has on-target activity under the conditions tested here.

A central finding of this work is the observation that PARG activity is dispensable for NAD^+^ depletion during parthanatos, but that ATP depletion requires (at least a small amount of) PARG activity (Fig. 6). This indicates that hyperactivation of PARP1 can deplete cellular NAD^+^ pools, without the need for PARG-dependent “recycling” of automodified PARP1. This data also suggests that the reduction in ATP levels is not a direct consequence of NAD^+^ loss, uncoupling NAD^+^ depletion from ATP depletion, as already observed previously in other settings ^[64,65]^. This indicates that even though NAD^+^ is required for adequate flux through metabolic pathways such as glycolysis, which could impact cellular ATP levels, the main mechanism by which PARP1 activation depletes ATP pools actually requires (at least a small amount of) PARG-dependent hydrolysis of PAR chains. Crucially, this finding also suggests that cell death by parthanatos correlates much better with ATP depletion than with NAD^+^ depletion, suggesting that ATP depletion is more likely to play a role in cell death execution than NAD^+^ depletion itself. As expected, this energetic collapse led to subsequent activation of AMPK, but whether this is an attempted cytoprotective response to ATP depletion or part of the core parthanatos programme required for cell death ^[66,67]^ is currently unclear.

Although several questions remain to be addressed regarding the detailed molecular mechanisms of parthanatos, and how these findings translate to models of parthanatos-dependent pathologies, this study demonstrates that, in addition to PARP1 hyperactivation, the hydrolysis of PAR chains by PARG is also essential for this cell death pathway, likely by promoting ATP depletion in response to PARP1 hyperactivity.

## Methods

### Cell culture

hTERT-RPE1 cells (ATCC: CRL-4000, referred to as RPE1 throughout the study) were grown in DMEM/F- 12 medium supplemented with 10% foetal bovine serum, 15 mM HEPES, and penicillin/streptomycin (100 U/mL / 100 μg/mL). A549 (ATCC: CRM-CCL-185) and HeLa (ATCC: CCL-2) cells were grown in DMEM high glucose medium (Thermo), supplemented with 10% foetal bovine serum and penicillin/streptomycin (100 U/mL / 100 μg/mL). All cultures were grown at 37 °C in a humidified atmosphere containing 5% CO_2_. Cells were seeded 24h prior to each experiment. The PARP1 KO hTERT-RPE1 cell line was generated in ^[68]^.

### Drug treatments

For MNNG treatment, cells were treated with fresh medium containing indicated doses of MNNG prepared immediately before each use. Olaparib and PDD0017273 were used at 10 µM unless otherwise stated and were added 1h before and during the MNNG treatment. Z-VAD-FMK was used at 50 µM and added 1h before and during the MNNG treatment. Staurosporine was used at 1 µM.

### XTT cell viability assays

RPE1 cells were seeded in 24-well plates at 2×10^4^ cells per well, incubated for 24h and treated with MNNG (above). After 72h of treatment, cells were washed with PBS and left in 200µL of XTT working solution for 3h. Cell Proliferation Kit II (XTT) (#11465015001 - Roche) was used. The working solution was prepared according to manufacturer’s instructions but diluted 8x in PBS. Supernatant was transferred to a 96-well plate and absorbance was read at 450nm and 750nm.

### Microscopy-based cell death assay and immunofluorescence microscopy

10^4^ cells were seeded on 96-well plates, treated the next day with MNNG and analysed 24h later. For cell death assays, cells were incubated in a PBS solution containing Hoechst 33342 (10 mg/mL) and propidium iodide (10 µg/mL) for 20 min under standard growth conditions. Cells were then imaged immediately and the images were analysed using StrataQuest software (TissueGnostics). Nuclear boundaries were detected based on Hoechst 33342 signal after background subtraction, and total propidium iodide signal was measured in each nucleus. Cells were considered PI-positive when signal reached a cut-off value, which was set above the fluorescence intensity of untreated samples. For immunofluorescence microscopy, cells were washed with PBS and fixed with PFA for 10 min, washed, permeabilized with PBS-Triton X-100 (0.2%) for 5 min, and blocked with 10% FBS in PBS. Primary antibody incubation was performed for 2 h at room temperature, followed by incubation with secondary antibody for 1 h. Cells were counterstained with DAPI, washed three times with PBS, mounted with 90% glycerol, stored at 4 °C, and imaged within 3 days. Images were analysed using StrataQuest software (TissueGnostics). Nuclear boundaries were detected based on DAPI signal after background subtraction, and total and mean ADP-ribose signal was measured in each nucleus. Microscopy images were acquired on a custom TissueFAXS iFluo system (TissueGnostics) mounted on an AxioObserver 7 microscope, as previously described ^[69]^.

### Western blotting

1-2 x 10^6^ cells per condition were seeded on 6-well plates and treated as appropriate. Adherent cells were washed with PBS and immediately lysed on-plate with 75–100 µL of incomplete Laemmli buffer (lacking β-mercaptoethanol/DTT and bromophenol blue), preheated to 100 °C. Cells were scraped, transferred to microtubes, and incubated at 100 °C for 10 min. After protein quantification using the Pierce™ BCA Protein Assay Kit (Thermo Fisher Scientific), sample volumes were normalized to equal protein concentrations, β-mercaptoethanol (5% v/v) and bromophenol blue were added, and 20-50 µg of protein per sample were loaded into SDS-PAGE gels. Proteins were transferred to nitrocellulose membranes at 100 V for 75–90 min, stained with Ponceau S, and blocked with 5% (w/v) BSA or 5% (w/v) milk in TBS-T. Primary antibody incubations were performed in blocking buffer; membranes were washed 3× in TBS-T, incubated with HRP-conjugated secondary antibodies, washed again and developed with ECL Prime (Amersham), and imaged on a ChemiDoc system (Bio-Rad).

### Antibodies

Antibodies used in this study are listed in Supplementary Table 1.

### Flow cytometry

1-2.5 x 10^6^ cells were seeded in 6-well plates, treated as appropriate the next day and collected 24h later. Culture supernatants were collected into 5-mL tubes. PBS washes were pooled with the supernatants to retain detached cells. Remaining adherent cells were trypsinized and combined with the pooled fractions. Cells were washed with PBS and resuspended in PBS containing propidium iodide (10 µg/mL) and immediately analysed by flow cytometry using a CytoFLEX LX Analyzer (Applied Biosystems) or an Attune Acoustic Focusing Cytometer (Beckman Coulter). A total of 10,000-30,000 events per condition were collected and analysed using FlowJo software. Cells were considered PI-positive when fluorescence reached a cut-off value, which was set above the fluorescence value of untreated samples.

### Guide RNA design and cloning

Custom single-guide RNAs (listed in Supplementary Table 2) were designed using Benchling ^[70]^. Oligonucleotides were phosphorylated at the 5′ end using T4 polynucleotide kinase (New England Biolabs) and annealed in a thermocycler ^[71]^. sgRNA inserts were ligated using T4 DNA ligase (New England Biolabs) into the eSpCas9(1.1) vector (Addgene #71814) linearized with BbsI, and transformed into NEB5α Escherichia coli (New England Biolabs) according to the manufacturer’s instructions. Plasmid DNA was isolated from bacterial cells using Monarch® Plasmid Miniprep (New England Biolabs) or QIAGEN Plasmid Midi (QIAGEN) kits and quantified using a NanoDrop 1000 spectrophotometer (Thermo Fisher Scientific). Constructs were verified by Sanger sequencing using a U6 promoter primer.

### CRISPR/Cas9 knockout

Vector transfection into RPE1 cells was performed using the Neon Transfection System (Thermo Fisher Scientific). For each condition, 3.2×10^5^ cells were electroporated using two pulses of 1150 V for 30 ms and subsequently plated into 6-well plates. CRISPR/Cas9 vectors were co-transfected with pCD2E vector (a kind gift of Keith Caldecott, Univ. Sussex) conferring G418 resistance. Transfected cells were selected by incubation with 1 mg/mL G418 for 5 days. In order to isolate knockout clones, transfected cell populations were seeded at low density (5-10 cells/mL) in 150mm dishes. Clones were left to grow for 1-2 weeks, and then transferred to separate wells in 96-well plates. After reaching confluency, clones were collected for western blotting and probed for the presence of the target protein. Genomic DNA was extracted from the chosen clone by standard high-salt/ethanol precipitation and the target region amplified by PCR

### Polymerase Chain Reaction (PCR)

PCR using the primers listed in Supplementary Table 2 was performed either on genomic DNA or using cDNA samples prepared in ^[72]^, using Q5 High-Fidelity DNA Polymerase (New England Biolabs) according to the manufacturer’s protocol, except that primer concentrations were decreased ten-fold.

### Sequencing

Sanger sequencing was performed using BigDye3.1™ Terminator v3.1 Cycle Sequencing Kit (Thermo Fisher Scientific) either in the Analytical Center of the Institute of Chemistry (USP) or in the Human Genome and Stem Cell Research Center (USP). Long-read Oxford Nanopore sequencing of PCR products was performed by PlasmidSaurus.

### Sequence analysis and isoform identification

Oxford Nanopore long reads were provided with adapters removed but without quality filtering and were used directly for downstream analyses. Transcript reconstruction was performed using StringTie2 (v. 2.2.1), with the -L parameter for long-read processing and the use of a reference annotation (GFF GRCh38). The assembly was descriptive in nature, aiming at the global identification of transcripts and isoforms present in the dataset, without subsequent use of the assembled transcripts as the primary unit of analysis. The sequences of the assembled transcripts were extracted from the GTF file using GFFread. For more in-depth analysis, raw reads were directly aligned to the human reference genome (GRCh38.p14) using minimap2 v2.11-r797 ^[73]^ with “-x splice” parameter. To more reliably discriminate transcripts derived from the *PARG* locus and the *PARGP1* pseudogene, only reads mapping primarily to either locus were considered and subsequently filtered based on exact matches to diagnostic single-nucleotide polymorphisms located in exon 9 (Supplementary Information). Reads containing the sequence *CATTTCCACGA**<u>C</u>**GAAATGCT**<u>A</u>**AGATGAAAT* were classified as *PARG*-derived, whereas reads containing *CATTTCCACGA**<u>T</u>**GAAATGCT**<u>C</u>**AGATGAAAT* were classified as *PARGP1*-derived. All subsequent analyses were restricted to these high-confidence read subsets. Identification and quantification of PARG isoforms were performed by direct detection of isoform-specific exon-exon junctions using exact sequence matching (100% identity, no mismatch tolerance). Splice junctions characteristic of each isoform were searched directly in the *PARG*-assigned reads using seqkit, including exon 1-2 junctions defining PARG111 (*GCCACCTCGCTTGTTTTCAAACA*) and PARG102 (*CATTGAGGCAGTTTTCAAACA* and *ATCTCTCTCAGGTTTTCAAAC*\*, which were combined), the exon 1-3 junction defining PARG99 (*GCCACCTCGCTTGTTTGGATAGTAAAG*), and the exon 1-8 junction defining PARG53 (*ACCTCGCTTGGTACTTGAAG*). The following splice junctions were searched based on the reported structures of PARG55/PARG60: (i) exon 1–4 junction “type PARG60” (*GTCCATCTCTCTCAGAAAAGAACA*); (ii) exon 1–4 junction “type PARG55” (*GTCCATCTCTCTCAGATAAGAAGT*); (iii) exon 4–6 junction “type PARG” (*ACTATTCGGAATGGTGAG*), (iv) exon 4-6 junction “type PARGP1” (*ACTGTTTGGAATGGTGAG*), and (v) exon 4-6 junction “type Meyer” ^[11]^ (*ACTATTTGGAATGGTGAG*). Reads containing different e1-e4 or e4-e6 junctions were combined in the graphs shown in Sup. Fig. 5 for simplicity. All searches were executed in an integrated manner within a single automated pipeline (script *search_juncs.sh*), which included pre-cleaning of FASTA files, filtering of reads containing the exon 9 SNPs, and subsequent detection of splicing junctions. Final counts were obtained directly from the resulting FASTA files, and the identifiers of reads positive for each motif were exported for downstream analyses. Isoform proportions were calculated based on the number of reads containing each diagnostic junction relative to the total number of PARG-assigned reads, excluding reads with none of the junctions found.

*This junction was not found in databases but was found in some of the transcripts encoding PARG102. This segment is untranslated in PARG102 and, therefore, did not affect predicted protein sequence.

### siRNA knockdown

SMARTPool (Dharmacon) siRNAs for the target proteins were used for most experiments. Custom siRNA sequences for PARG53 were designed by Thermo Fisher Scientific. siRNA sequences are listed in Supplementary Table 3. For transfections, 1.5×10^4^ cells were seeded in 6-well plates, and 15 pmol siRNA were mixed with Lipofectamine RNAiMAX (Thermo) following the manufacturer’s protocol, using both the forward and reverse transfection protocols on the same cells. Cells were harvested 48h after the second transfection for western blotting.

### Nucleotide extraction and HPLC

Nucleotide extraction, HPLC and protein quantification were performed as described in ^[56]^. In short, cells were seeded in 10 cm dishes and, after treatment, cells were fixed and permeabilized in methanol supplemented with 1 mM deferoxamine mesylate. Cells were immediately scraped and samples transferred to microtubes and mixed until completely dissolved. After centrifugation, the supernatant was vacuum-dried and stored at -80°C, while the precipitate was stored at -20°C for protein quantification and western blotting. For HPLC column specifications and separation method used, please refer to the aforementioned paper.

### Quantification and statistical analysis

Statistical analyses were performed using GraphPad Prism 8.0.1 software. Details on statistical tests, sample sizes, and replicate numbers are reported in the figure legends. Statistical significance was defined as p < 0.05 unless otherwise indicated.

## Acknowledgements

We would like to thank Célia Ludio Braga for technical assistance in the NCH lab, the Chemistry Institutés “Central Analítica” for assistance with the HPLC experiments and Prof. Henning Ulrich for use of the flow cytometers. This work was funded by FAPESP grants 2018/18007-5 and 2020/05317-6, as well as Wellcome Trust Career Development Award 314035/Z/24/Z to NCH. JCS is the recipient of a CNPq Research Fellowship (305274/2024-4). MHGM and NCSP acknowledge funding from FAPESP (2013/07937-8 and 2024/07910-7, respectively). We gratefully acknowledge stipends from FAPESP to RDM (2020/02701-0) and BK (2024/03685-9), from CNPq to PDM and RTN, and from CAPES to PFV and ACS. The microscope used in this study was funded by FAPESP grant 2019/06039-2 (to NCH).

## Author contributions

Conceptualisation and study design: RDM, NCH. Method development and validation: RDM, BK, PDM, PFV, ACS, RTN, FMP. Data acquisition: RDM, BK, PDM, PFV, ACS. Data analysis: RDM, BK, PDM, PFV, ACS, RTN, NCH. Funding acquisition and supervision: MHGM, JCS, NCSP, NCH. Manuscript writing: RDM, NCH. Manuscript revision: all authors.

## SUPPLEMENTARY DATA

**Supplementary Figure 1:**
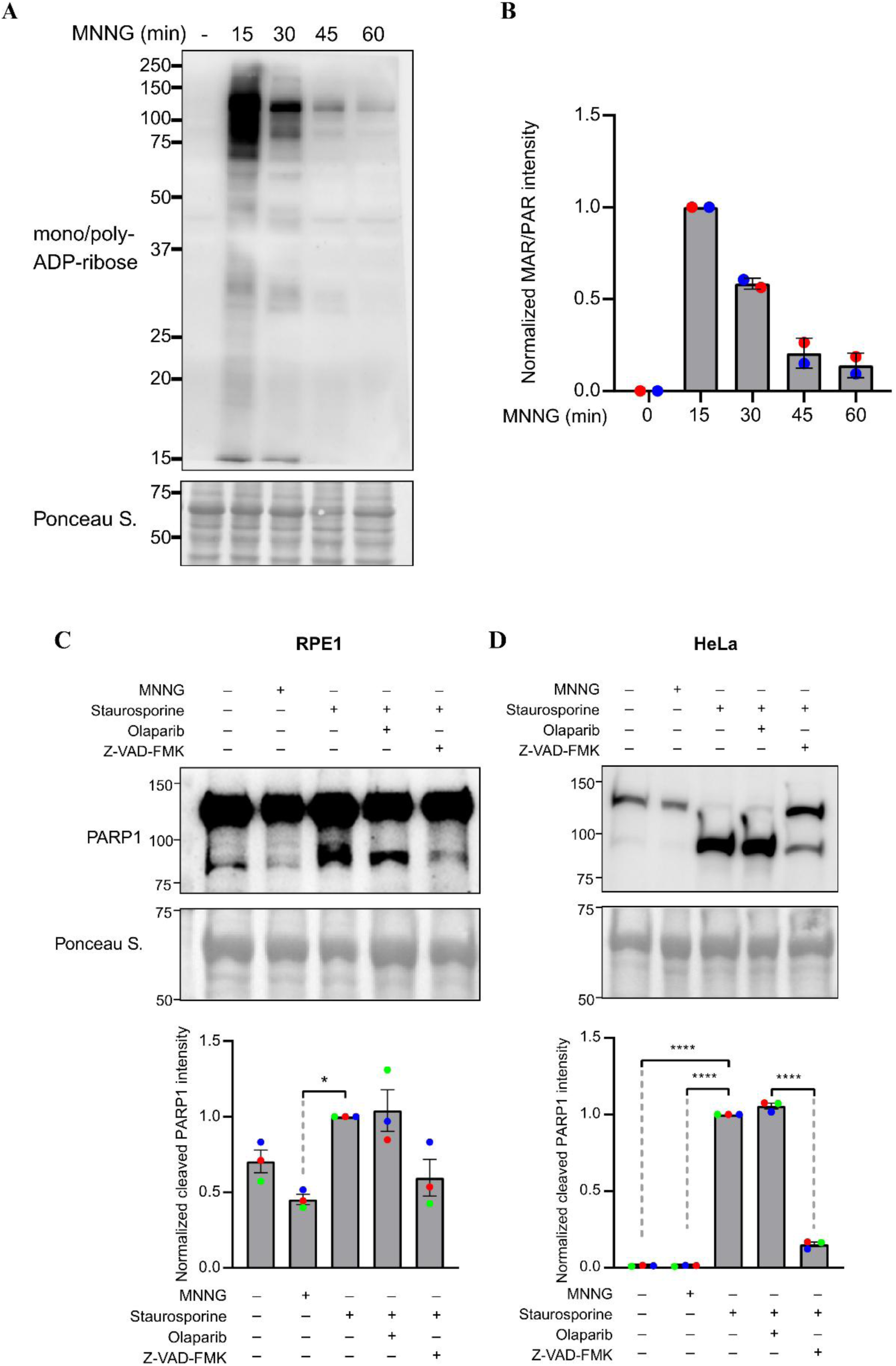
MNNG-induced parthanatos involves a wave of ADP-ribosylation and is a non-apoptotic form of cell death. **A)** Representative western blot and **B**) quantification of total protein ADP-ribosylation after treatment of RPE1 cells with 250uM MNNG for the indicated time periods. **(C and D)** Representative western blots for PARP1 protein (top) and quantification of PARP1 cleavage (bottom) in RPE1 cells (C) or HeLa cells (D) treated with MNNG (250 µM for RPE1 cells and 2.5 mM for HeLa cells), olaparib (10 µM), the apoptosis-inducing agent staurosporine (1 µM) and/or the pan-caspase inhibitor z-VAD-FMK (50 µM), as indicated. Cells were harvested after 6h of MNNG or staurosporine treatment. Graphs depict mean and SEM of three independent experiments, normalized to the staurosporine-treated sample of each replicate.

**Supplementary Figure 2:**
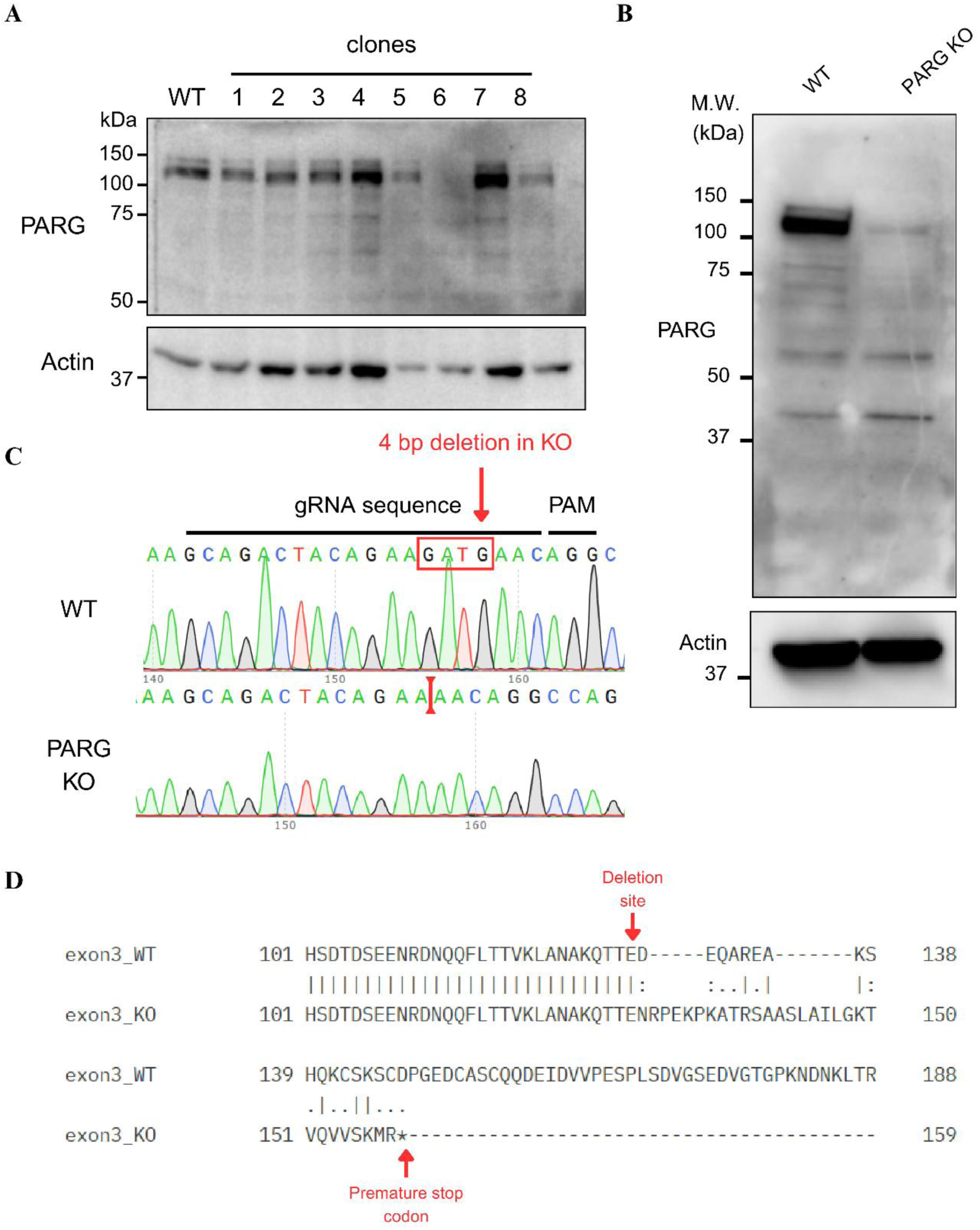
Generation of a PARG knockout cell line. **A)** Screening of *PARG* knockout clones. Individual clones were isolated from a population of cells transfected with the CRISPR/Cas9 vector targeting *PARG* (not all are shown). Clone #6 was established as our PARG KO cell line for all further experiments in this work. **B)** anti-PARG immunoblot comparing KO clone #6 with RPE1 WT controls. **C)** Sanger sequencing of PCR products from the Cas9 target region in the *PARG* gene showing the homozygous 4nt deletion in clone #6. **D)** Global pairwise alignment of translated WT and KO sequences, indicating a premature stop codon formed as a result of the 4 nucleotide deletion.

**Supplementary Figure 3:**
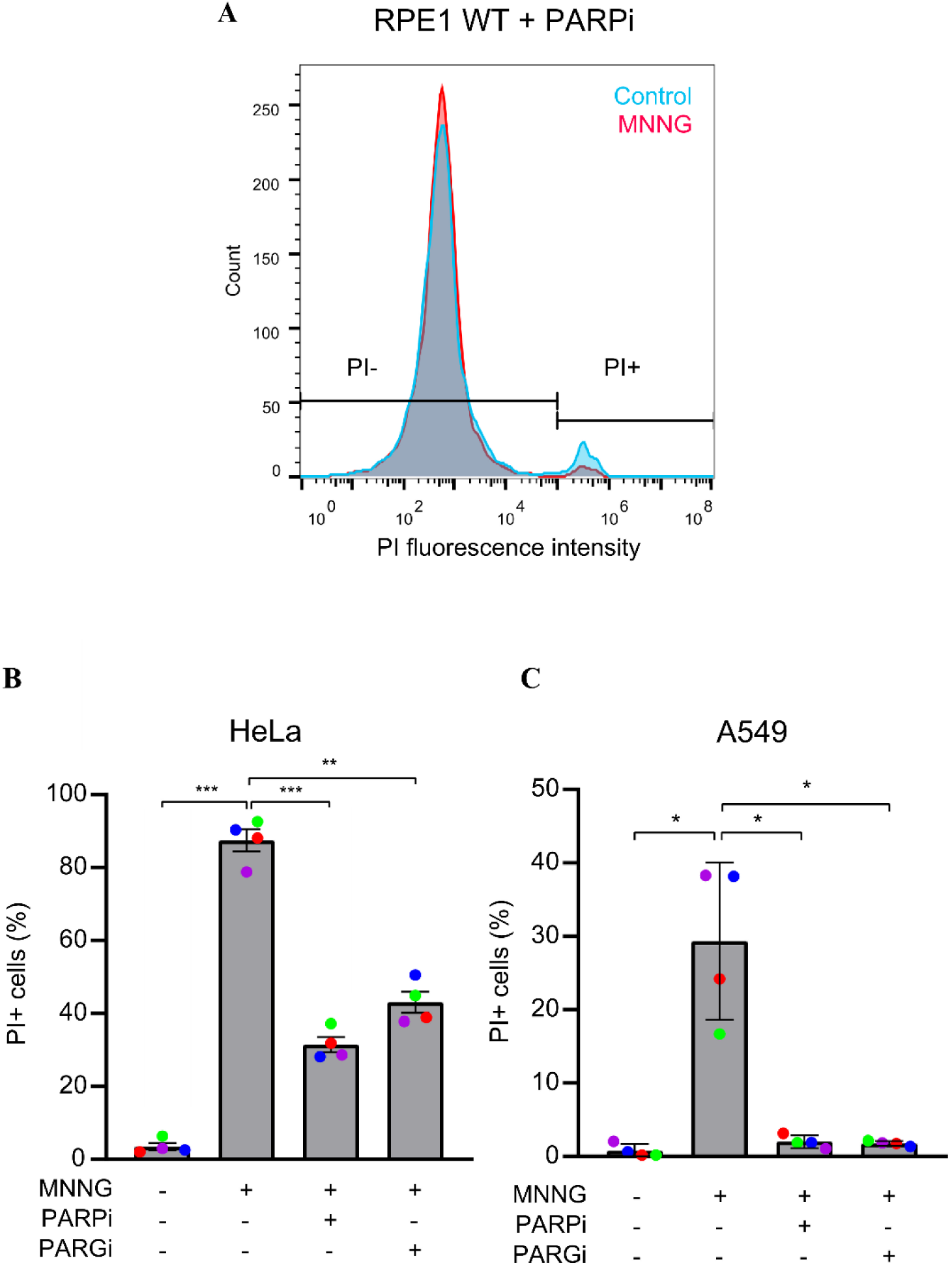
Both PARP1/2 and PARG inhibition prevent parthanatos in human cell lines. **(A)** Representative histogram overlay of PI fluorescence intensity in WT RPE cells treated for 24h with olaparib only (blue) or with olaparib plus MNNG (red) (related to Figure 3A-B). **(B and C)** Quantification of PI-positive cells after treatment of HeLa cells (B) or A549 cells (C) with MNNG (at 500 µM for A549 and 2.5 mM for HeLa cells), 10 µM olaparib and/or 10 µM PARG inhibitor. Data were analysed using one-way ANOVA with Tukey’s test (ns = not significant, * = p < 0.05, ** = p < 0.005, *** = p < 0.0005).

**Supplementary Figure 4:**
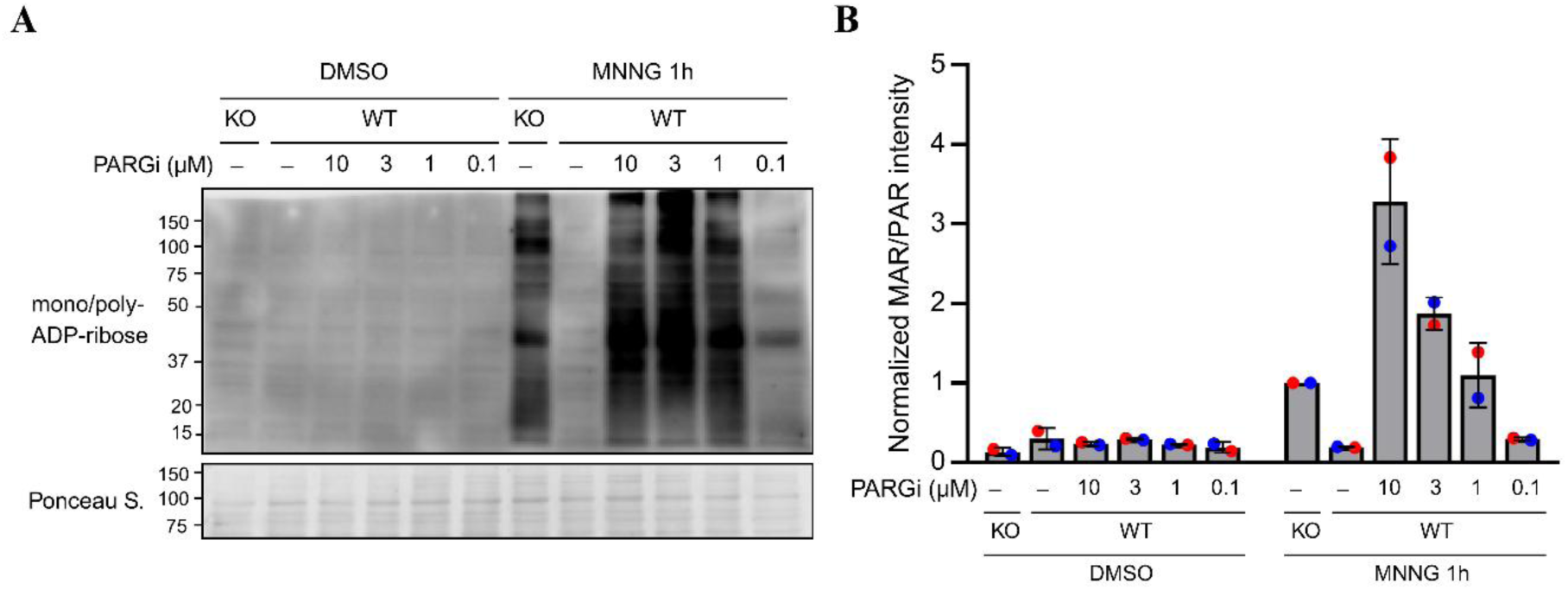
Effect of lower PARG inhibitor concentrations on PAR hydrolysis. **(A)** Representative western blot and **(B)** quantification of mono/poly-ADP-ribosylation in PARG KO or WT RPE1 cells treated for 1h with DMSO control or MNNG, as well as PARG inhibitor at indicated dose

**Supplementary Figure 5:**
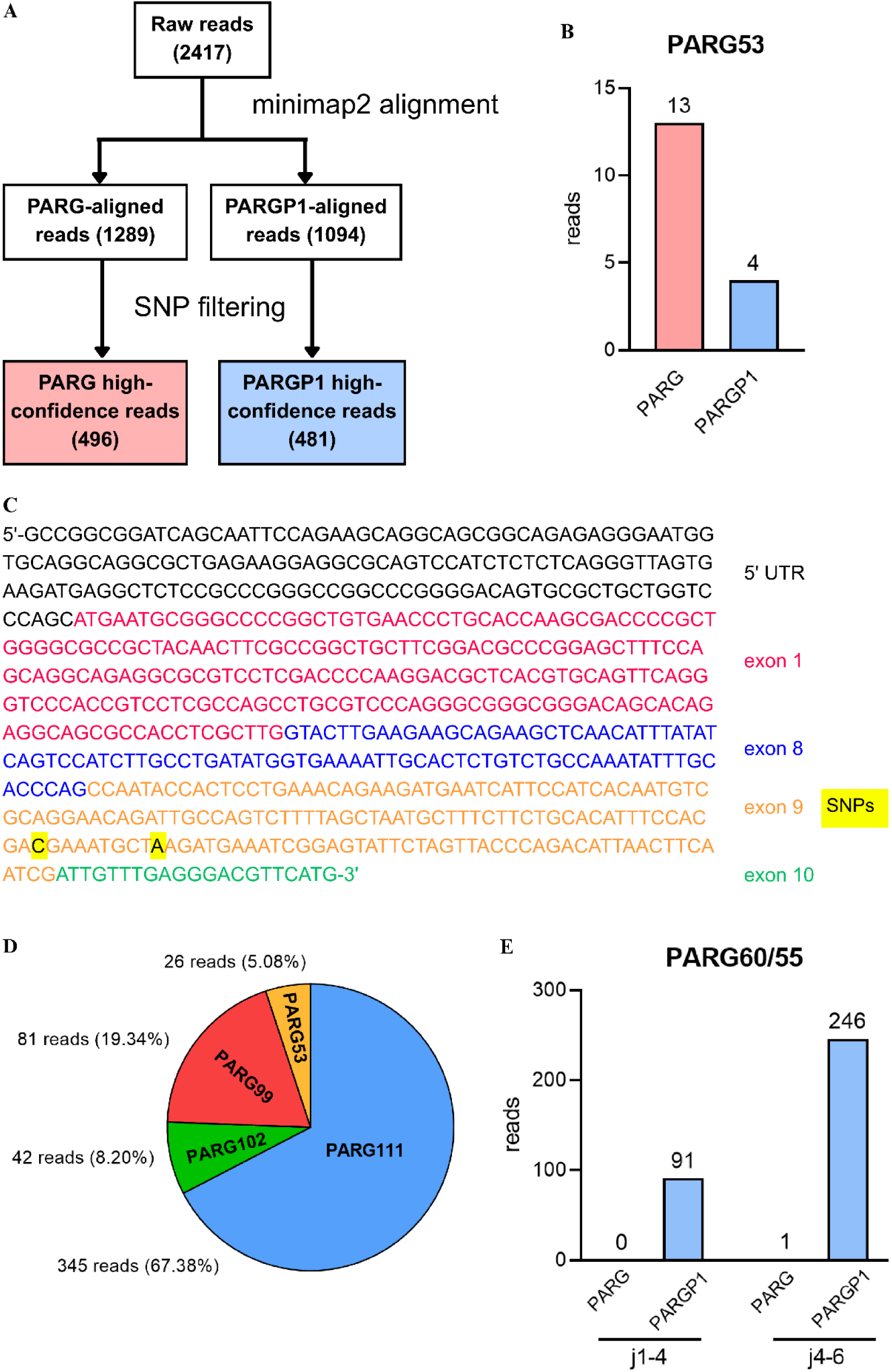
Identification of PARG isoforms. **A)** Methodology used for separating *PARG*- and *PARGP1*-derived sequences. Reads were classified based on their primary alignment. Diagnostic SNPs in exon 9 were used for verification and filtering. **B)** Number of reads originated from *PARG* and *PARGP1* containing the PARG53 junction (exon1-exon8). **C)** Example of a raw read encoding the PARG53 isoform, with diagnostic SNPs highlighted, validating its identification as a *PARG* sequence. **D)** Percentage of reads encoding each isoform, identified by searching their respective splice junctions. Reads containing none of the junctions were not included in the total. **E)** Number of reads originated from *PARG* and *PARGP1* containing the PARG60/55 junctions (exon1-exon4 and exon4-exon6).

**Supplementary Figure 6:**
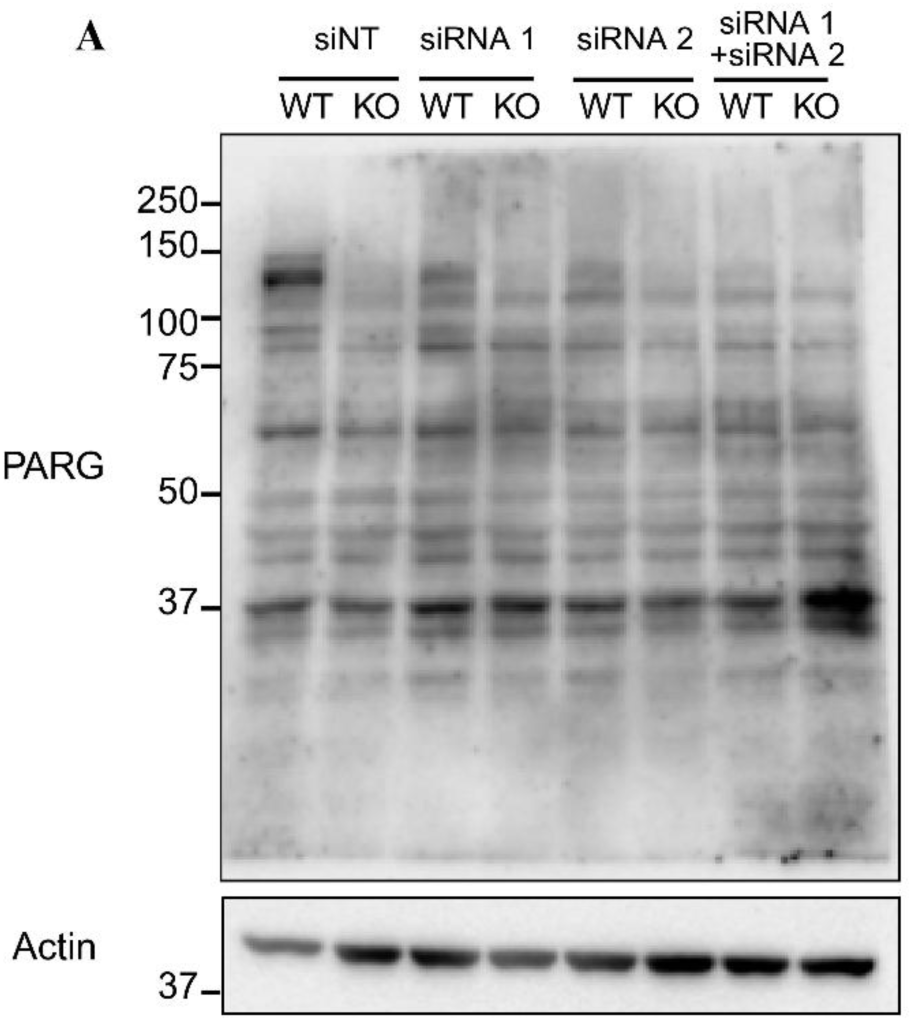
Attempts at detecting PARG53 at the protein level via siRNA knockdown. **A)** Anti-PARG blot showing knockdown of PARG using two different custom siRNAs specifically designed to target the exon 1-8 junction in PARG53.

**Supplementary Figure 7:**
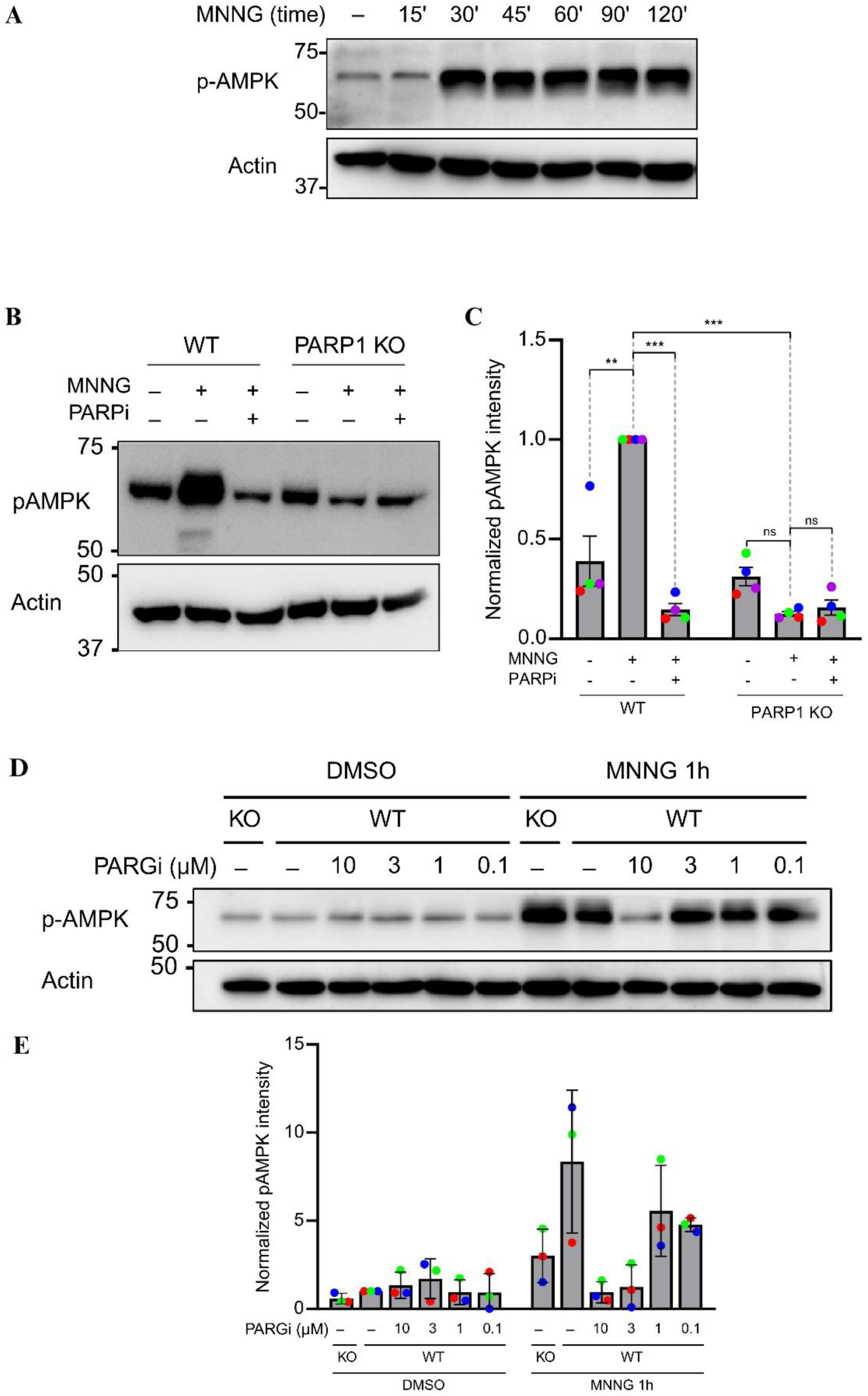
Parthanatos-induced AMPK activation requires PARP1 and PARG activities. **A)** Time course of AMPK Thr172 phosphorylation in RPE1 cells treated with MNNG. **B)** Representative western blot and **(C)** quantification of AMPK Thr172 phosphorylation in WT or PARP1 KO RPE1 cells treated for 1h with MNNG or olaparib, as indicated. **(D)** AMPK phosphorylation in PARG KO or WT RPE1 cells after 1h of treatment with DMSO control or MNNG, as well as indicated doses of PARG inhibitor.

**Supplementary Figure 8:**
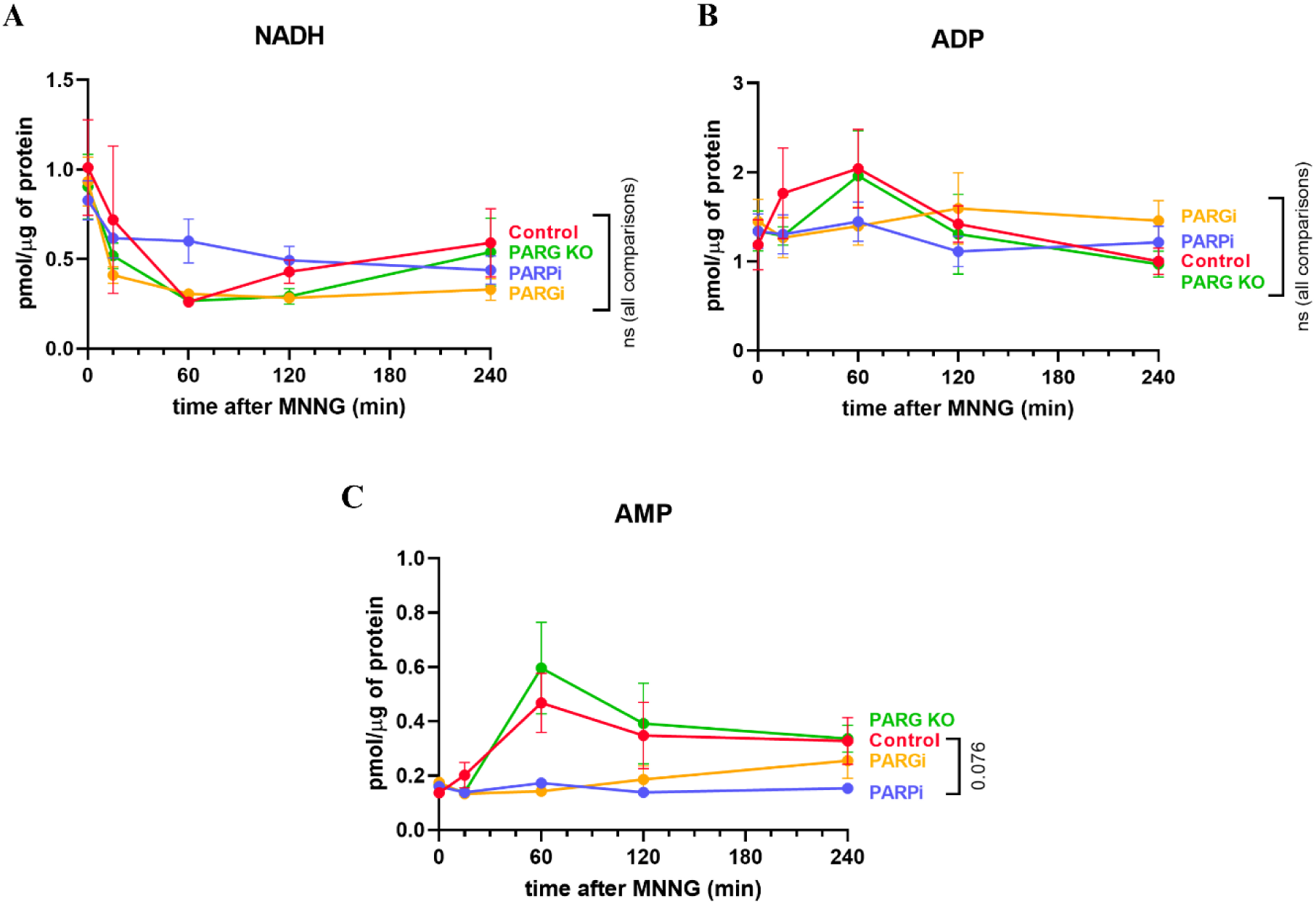
Nucleotide quantification by HPLC of NADH, ADP and AMP. Quantification of NADH (A), ADP (B) and AMP (C) in samples from Fig. 6 B-C.

**Supplementary Table 1.** List of primary antibodies used in this study.

| Target | Clone | Host species | Application | Supplier | Catalogue no. |
| --- | --- | --- | --- | --- | --- |
| Pan-ADP-ribose | – | Rabbit | WB, IF | Millipore | MABE1016 |
| Mono-ADP-ribose | AbD43647 | Human (recombinant Fab) | WB | Bio-Rad | TZA020P |
| H3-S10-ADP-ribose | AbD33644 | Rabbit | WB | Bio-Rad | HCA357 |
| PARP1 | – | Rabbit | WB | Abcam | ab110915 |
| PARP1 (cleaved) | – | Rabbit | WB | Cell Signaling Technology | #9542 |
| PARG | D8B10 | Mouse | WB | Millipore | MABS61 |
| Phospho-AMPKα (Thr172) | 40H9 | Rabbit | WB | Cell Signaling Technology | #2535 |
| Actin | C4 | Mouse | WB | Millipore | MAB1501 |
| α-Tubulin | – | Mouse | WB | Abcam | ab18251 |

**Supplementary Table 2.**
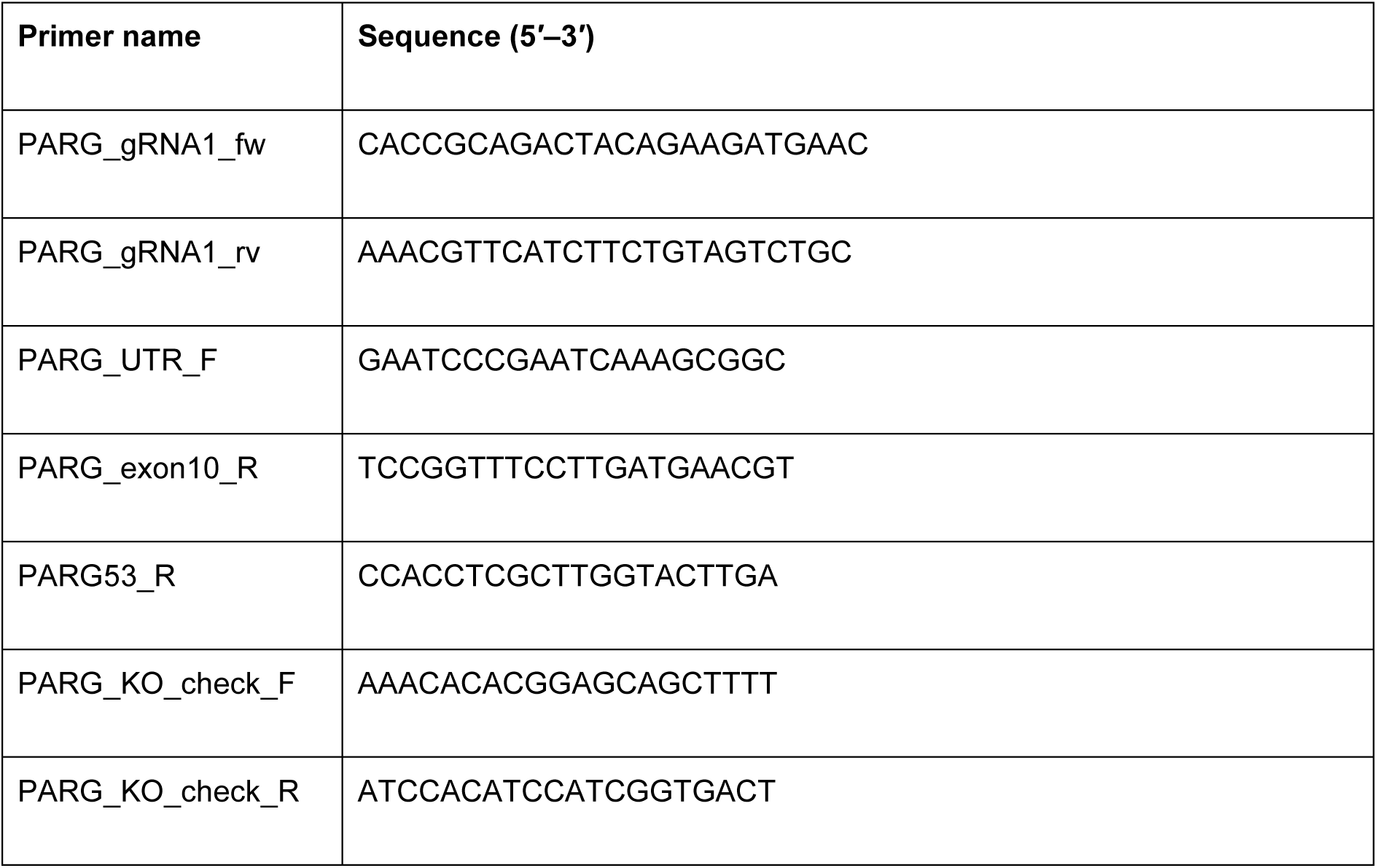
List of oligonucleotides used in this study.

**Supplementary table 3.**
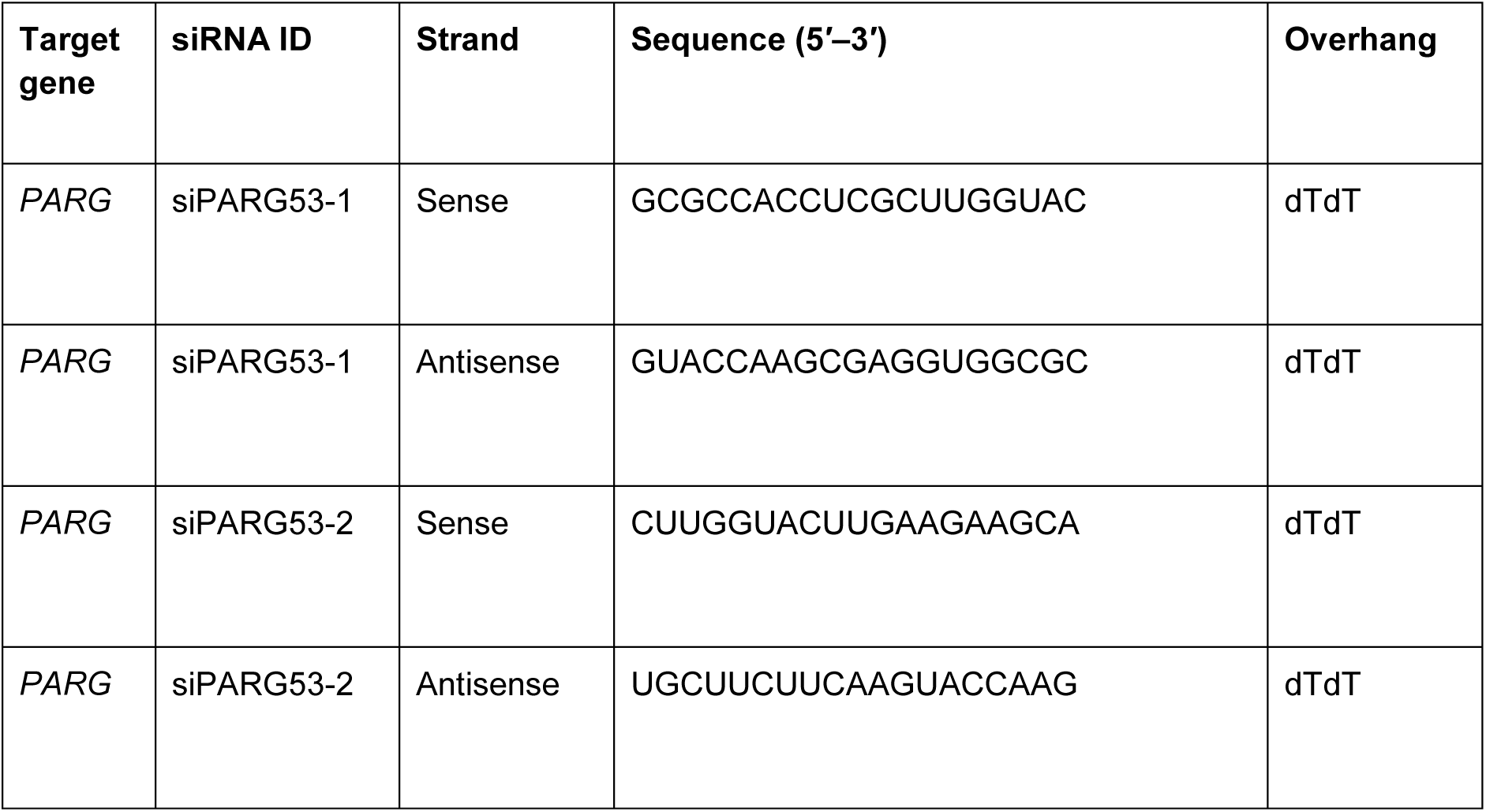
List of siRNAs used in this study.

## Supplementary Information describing the analysis of Oxford Nanopore sequencing

As discussed in the main text, PCR on RPE1 cDNA using a forward primer in the 5′ UTR and a reverse primer within exon 10 of the *PARG* gene (Fig. 5B) yielded a number of products of varying sizes (Fig. 5C). This material was sent for Oxford Nanopore sequencing, which generated 2417 raw reads (Supplementary Data File 1) that were used for all analyses described below.

First, we sought to determine the splicing pattern of these reads, so StringTie was used to assemble these reads into a collection of different transcripts (Table 1 and Supplementary Data File 2). The consensus sequences for each of these transcripts were then searched using BLAST, which identified that several of these consensus transcripts were more similar to the *PARGP1* pseudogene than to the *PARG* gene itself (Table 1). We confirmed that the primer sequences we used for the PCR are present within the *PARGP1* pseudogene, indicating that their amplification was specific (data not shown). This data indicates that the *PARGP1* pseudogene is transcribed, that its transcripts are spliced, and that a substantial proportion of our 2417 raw reads originated from the *PARGP1* pseudogene.

To more accurately distinguish the genomic origin of each read, we used MiniMap to map each individual read to the human genome. For 1289 raw reads (53.3% of the total) the primary alignment was to the *PARG* gene, whereas 1094 raw reads (45.3% of the total) aligned preferentially to the *PARGP1* pseudogene (Sup. Fig. 5A). The remaining 34 reads (1.4%) could not be determined and were discarded. The *PARG* gene and the *PARGP1* pseudogene are highly similar, but several single nucleotide polymorphisms (SNPs) scattered throughout the sequence allow clear distinction between them. We focused on a pair of closely spaced SNPs in exon 9 in which the *PARG* sequence is CGA**<u>C</u>**GAAATGCT**<u>A</u>**AGA, whereas the *PARGP1* sequence is CGA**<u>T</u>**GAAATGCT**<u>C</u>**AGA. All raw reads were searched for the presence of either of these sequences, which allowed us to generate a high-confidence set of 496 reads that aligned preferentially to the *PARG* gene and also contained the exon 9 SNPs from the *PARG* gene, as well as a high-confidence set of 481 reads that aligned to the *PARGP1* pseudogene and also contained the exon 9 SNPs from the *PARGP1* pseudogene (Sup. Fig. 5A). Over 90% of the reads that were filtered out in this step did not contain any exon 9 sequence and could not be confidently assigned, while less than 3% of the reads had discordant alignment/SNP filtering results (data not shown).

Using this set of high-confidence raw reads, we searched for the identified exon1-exon8 junction that characterizes the novel transcript predicted to encode PARG53 (transcript 5 - Table 1), to determine its genomic origin. We found 13 high-confidence reads that originated from the *PARG* gene and contained the exon1-exon8 junction, and 4 high-confidence reads that originated from the *PARGP1* pseudogene and contained this junction (Sup. Fig. 5B). The full raw sequence of one such read from the *PARG* gene is shown in Sup. Fig. 5C. This data confirms that mRNA transcribed from the *PARG* gene can be spliced from exon1 into exon 8, to generate a transcript predicted to encode PARG53, and indicates that the same splicing event can also occur in transcripts originating from the *PARGP1* pseudogene.

Next, we used the same strategy to determine the genomic origin of reads that contained the exon1-exon4 junction and the exon4-exon6 junction, that are characteristic of the PARG60 and PARG55 isoforms ^[8,11]^. While none of the high-confidence reads from the *PARG* gene contained the e1-e4 junction, 91 out of 481 (18.9%) of the high-confidence reads from the *PARGP1* pseudogene contained this junction (Sup. Fig. 5D). Similarly, only one of the 496 reads from the *PARG* gene (0,2%), but 246 out of 481 reads from *PARGP1* pseudogene (51,1%) contained the exon4-exon6 junction (Sup. Fig. 5D). Pseudogenic transcripts cannot encode full-length PARG55 or PARG60, as the *PARGP1* pseudogene only contains sequences up to exon 13 out of the 18 exons in the *PARG* gene (data not shown). Additionally, the most frequently detected splicing structure from pseudogenic transcripts (transcript P1 - Table 1), contained an additional exon between exon 7 and exon 8, changing the reading frame (data not shown). Collectively, these results indicate that the exon1-exon4-exon6 splicing structure that characterizes the reported PARG55/PARG60-encoding transcripts are likely to have originated from PCR amplification of transcripts from the *PARGP1* pseudogene, suggesting that PARG55 and PARG60 are not true PARG isoforms.

To gain some approximate information about the relative abundance of transcripts encoding PARG isoforms, we quantified the number of raw reads that contain the characteristic splice junctions for transcripts encoding PARG111 (junction e1-e2), PARG102 (5’UTR-e2 junctions, of which there are two), PARG99 (junction 5’UTR-e3) and PARG53 (junction e1-e8) (Fig. 5B). This search was limited to the set of 1289 raw reads that aligned to the *PARG* gene (Sup. Fig. 5A), and yielded 512 raw reads that could be confidently assigned to either of the isoforms. 355 of these 521 reads (67.38%) contained the splice junction for the PARG111-encoding transcript, while the junctions for transcripts encoding PARG102 were found in 42 reads (8.2%). Characteristic junctions for PARG99 were found in 99 reads (19.34%) and 26 reads contained the e1-e8 junction for PARG53 (5.08%) (Sup. Fig. 5E). Although transcript abundance does not necessarily correlate with protein abundance, these data suggest that PARG111 is the most abundant PARG isoform, as expected, followed by PARG99, PARG102 and PARG53.

